# Past and future phenology changes of zoonotic vector-borne diseases under climate and land-use change

**DOI:** 10.64898/2026.06.07.730142

**Authors:** Valén Holle, Raphaëlle Klitting, Nadja Kabisch, Damaris Zurell

## Abstract

Environmental changes are reshaping the distribution and seasonal dynamics of vector-borne diseases, with important implications for public health. Tick-borne encephalitis virus (TBEV) and West Nile virus (WNV) cause growing concern in Europe, with rising case numbers and ever-expanding circulation areas. The transmission risk of TBEV and WNV follows characteristic seasonal patterns, driven largely by weather-dependent activity of their arthropod vectors. The relative roles of climate and land-use change on the seasonal dynamics and spread of these diseases and their vectors remain, however, poorly quantified. Here, we assess the spread and phenology of TBEV and WNV in response to historical and future climate and land-use changes across Europe. We developed spatiotemporal species distribution models (SDMs) for the viruses and their primary vector species, generating monthly environmental suitability predictions from the 1970s to 2050s. Virus models incorporated vector suitability as a nested predictor to capture the dependence of virus occurrence on vector presence. To disentangle drivers of observed changes, we applied counterfactual historical simulations, attributing shifts in seasonal transmission risk to climate or land-use changes. Historical attribution results show that land-use changes mainly affected absolute vector suitability, whereas climatic changes drove shifts in seasonal transmission risk. Transmission risk is projected to rise continent-wide for both TBEV and WNV over the coming decades. Further, TBEV is projected to undergo pronounced phenological shifts, with a dominant spring peak and a delayed autumn peak extending into October. Prolonged seasonal transmission windows are projected to create hotspots that both intensify and expand across large regions. Taken together, our findings underscore the need for coordinated transnational efforts to manage the projected health burden of TBEV and WNV across Europe, and support upstream prevention by providing climate-informed guidance on intervention timing and spatial prioritisation.

## Introduction

Accelerating environmental change is reshaping the ecology of infectious diseases (McElwee et al., 2024). As climate warming and habitat alteration shift the distribution of species, the interactions among pathogens, hosts, and vectors are also changing, leading to a redistribution of zoonotic vector-borne diseases and shifts in their seasonal emergence (Franklinos et al., 2019). Notably, tick-borne encephalitis virus (TBEV) and West Nile virus (WNV), both members of the genus *Orthoflavivirus* within the family *Flaviviridae*, are posing an increasing public health challenge in many parts of Europe (Barrett, 2018; Riccardi et al., 2019). The disease burden of TBEV is particularly high in endemic regions, including parts of Central and Eastern Europe, Scandinavia, and the Baltics (European Centre for Disease Prevention and Control, 2024), with a notable rise and northwestward expansion of TBEV infections over the past decade (Heuverswyn et al., 2023). The most recent data on annual WNV infections indicate, however, a more spatially restricted distribution with regional spikes in Southern and Eastern Europe, and notable outbreaks in Greece, Romania, and Italy (European Centre for Disease Prevention and Control, 2021). Similar to TBEV, the virus is continuing to spread, and new infection areas are emerging in previously unaffected regions such as Germany and Slovakia (European Centre for Disease Prevention and Control, 2021).

Global change drivers such as climate and land-use change can directly and indirectly affect such disease spreading through the complex biotic interplay of and transmission between hosts, vectors, and pathogens. In Europe, TBEV is primarily transmitted through the bite of the infected tick species *Ixodes ricinus* (Linnaeus, 1758), which serves as the main vector, acquiring the virus from infected small mammals and passing it on to humans and other hosts (Dagostin et al., 2024). For WNV, the transmission cycle involves the mosquito species *Culex pipiens* (Linnaeus, 1758) as the primary vector in Europe, acquiring the virus from infected wild birds and transmitting it to humans and other mammals, which act as dead-end hosts (Tran et al., 2014). Thereby, climate acts as a limiting factor for the distribution of both vectors and viruses, while land use primarily affects the availability of suitable habitat and thus the spatial distribution of vectors, with more indirect effects on virus transmission. For example, the proportion of forested area has been found to impact TBEV incidence in Europe, likely by increasing the encounter rate of tick vectors and hosts through the provision of suitable habitats (Dagostin et al., 2023). Similarly, urban and agricultural areas have been identified as contributing to WNV incidence by providing suitable habitats for the *Culex pipiens* mosquitoes, which thrive in close association with human-modified environments (Kilpatrick, 2011).

While climate and land-use change are increasingly considered in the studies of disease spread (Klitting et al., 2022; Lippi et al., 2023), their influence on disease phenology, i.e., changes in seasonal timing, is often overlooked. A particular characteristic of both TBEV and WNV is their seasonal emergence patterns, driven by the weather-dependent activity of their arthropod vectors. In recent years, the temporal distribution of reported TBEV infections in Europe has consistently shown two peaks, typically occurring in July and November (European Centre for Disease Prevention and Control, 2024). These seasonal dynamics reflect the close coupling between vector activity, virus transmission, and climatic conditions. Activity of *Ixodes ricinus* has been found to be positively correlated with temperature and influenced by other climatic factors such as higher relative humidity, which is particularly important for the survival of nymph stages that are highly susceptible to dehydration (Schulz et al., 2014; Zając et al., 2021). Higher humidity has also been shown to strongly affect TBEV prevalence in ticks (Lamsal et al., 2023). WNV cases predominantly occur between July and October, with infections peaking in August (European Centre for Disease Prevention and Control, 2021). Elevated temperatures and rainfall fluctuations have been identified as key factors sustaining local endemic WNV circulation in Europe and facilitating the establishment of the virus in new areas (Paz, 2015). Higher temperatures accelerate viral replication and promote the growth of the vector species *Culex pipiens,* while rainfall fluctuations have a dual effect: increased rainfall expands the surface area of standing water, which is crucial for larval development, while drought conditions concentrate organic material, providing essential nutrients for mosquito breeding (Paz, 2015). Predicting how climate-driven shifts in the seasonal timing of vector and virus activity will influence future disease phenology supports anticipating periods of elevated transmission risk and informing public health strategies for TBEV and WNV in Europe.

Despite the recognised importance of seasonal dynamics in disease transmission, there is a notable gap in our predictive understanding of TBEV and WNV spread and phenology in Europe, particularly regarding how these patterns have evolved historically and how they may shift under future environmental change scenarios. Previous assessments of disease seasonality have primarily relied on field surveys, often covering only limited time frames (Borde et al., 2019; Reynolds et al., 2022; Siperstein et al., 2023). Scaling up to broad spatial and temporal scales requires a comprehensive spatiotemporal modelling framework and long-term analyses that integrate both vector and virus distributions, enabling a deeper understanding of how the phenology of TBEV and WNV responds to changing environmental conditions in Europe. Correlative species distribution models (SDMs) have been widely used to understand disease spread (Aliaga-Samanez et al., 2024; Fischer et al., 2025) and to project future risk hotspots (Aldwekat et al., 2025; Klitting et al., 2022). However, while these models have provided valuable insights into the potential distribution of vectors and viruses, most studies have treated vectors and pathogens separately and have often been geographically limited to specific countries (McDonough & Holloway, 2020; Mezhzherin et al., 2024; Tytar et al., 2024; Uusitalo et al., 2022), with comparably few studies addressing continent-wide patterns across Europe (Alkishe et al., 2017; Erazo et al., 2024; Walter et al., 2020) and without explicit consideration of phenology shifts. To address this gap, we here suggest an SDM-based modelling framework that explicitly integrates vector and virus models in a nested design. This approach allows us to capture the dependency of virus transmission on vector presence by using predicted vector habitat suitability as a covariate in the virus SDMs, providing a more ecologically realistic framework for assessing spatial and temporal disease risk. By explicitly considering temporal components of the niche (Zurell et al., 2024), the SDM-based modelling framework allows us to account for disease phenology and elucidates how interannual and seasonal environmental variations affect the habitat suitability for both vectors and viruses throughout the year. We additionally combine this approach with long-term time series analyses within a detection and attribution framework (Gonzalez et al., 2023; Thomas et al., 2026), improving our ability to assess how climate and land-use change have historically shaped disease distribution and transmission patterns as well as phenology. Building on this improved understanding, we can further evaluate whether ongoing and future changes in climate and land use are likely to reinforce these trends, driving the expansion and shifting of endemic regions in Europe and altering disease transmission patterns (Gottdenker et al., 2014; Kilpatrick & Randolph, 2012; Lafferty, 2009; Ostfeld, 2009), potentially placing an increasing portion of the population at risk over longer time periods (Dagostin et al., 2024; Farooq et al., 2023). Overall, the framework provides a basis for anticipating spatiotemporal patterns of disease risk and supports more effective public health strategies at the regional to continental scale, including early warning systems, targeted surveillance, and adaptive public health strategies across regional to continental scales.

In this study, we aim to assess the historical and potential future impacts of climate and land-use changes on the disease spreading and phenology of TBEV and WNV in Europe. Using spatiotemporal SDMs, we generate monthly environmental suitability predictions for the main vector species and the associated viruses. We first quantify historical shifts in spatial distribution and seasonal dynamics between 1970 and 2019 and attribute these changes to climate and land-use drivers using a counterfactual-based impact attribution framework (Frieler et al., 2024; Schrodt et al., 2025; Thomas et al., 2026). We then project future disease spread and phenological changes for 2030-2059 under climate and land-use scenarios. Based on these analyses, we aim to address three main research questions:

1. How are risk distribution patterns changing from the historical to future periods?
2. How are phenological patterns in risk intensity changing from historical to future periods, and how do these patterns vary across space?
3. How are phenological patterns in risk duration changing in the future?

Together, this framework allows us to disentangle and quantify how climate and land-use change have shaped and continue to shape the spatial distribution and seasonal dynamics of infectious disease risk under global change.

## Materials and Methods

We built spatiotemporal SDMs for both the viruses and their main vector species across Europe using an ensemble modelling approach. All analyses were conducted at a 0.5° spatial resolution, which represented the finest common resolution available for all input datasets. First, we modelled the spatiotemporal distribution of the vector species, incorporating monthly climate variables and annual land-use data as predictors. Second, we modelled the spatiotemporal distributions of the viruses using a nested SDM framework, in which the predicted monthly habitat suitability of the respective vector species was included as a predictor variable in the virus models alongside monthly climate data. Third, to attribute observed changes in the distributions and phenology of both vectors and viruses, we predicted historical distributions under counterfactual climate and counterfactual land use scenarios. Finally, we predicted future changes in distributions and phenology under multiple climate and land-use scenarios. We provide an ODMAP protocol (Zurell, Franklin, et al., 2020; Supplementary Table S1), where all data preparation and modelling steps are detailed.

### Species data

Spatially explicit occurrence data for the primary vector species *Ixodes ricinus* and *Culex pipiens* were obtained from GBIF (Global Biodiversity Information Facility; https://www.gbif.org/; GBIF.org, 2025a, 2025b) and VectorMap (http://vectormap.si.edu). As an initial filtering step, records were restricted to a European bounding box (31°W, 40°E, 34°N, 72°N) and to the period 1970-2019, selected to reflect improved data consistency and reliability from 1970 onwards and the availability of observed environmental data up to 2019. Additionally, occurrence data were cleaned (Zizka et al., 2019) and aggregated to a 0.5° spatial and monthly temporal resolution, with duplicate records within grid cells and calendar months of a given year removed.

Since most SDM algorithms require background data in addition to presence data to identify areas with unsuitable environmental conditions, we generated background points for *Ixodes ricinus* and *Culex pipiens* by randomly selecting locations within species-specified buffer distances from the presence points, excluding cells containing observed presence records (150 km for *Ixodes ricinus*; 100 km for *Culex pipiens*) (Barbet-Massin et al., 2012). Background points were temporally matched to occurrences within the same month of a given year. Following established recommendations, we aimed for a presence-to-background ratio of 1:10 for regression-based algorithms and 1:1 for machine-learning algorithms (Barbet-Massin et al., 2012). Lastly, to avoid spatial autocorrelation, the species’ monthly presence and background data were thinned using a 50 km threshold (Aiello-Lammens et al., 2015). Supplementary Figure S1 shows a map of all presence points for the vectors and the generated background points.

### Infection data

TBEV and WNV human infection data for Europe were accessed through TESSy (The European Surveillance System) and released by the ECDC (European Centre for Disease Prevention and Control). The data included information on locally acquired confirmed cases, aggregated on the NUTS (Nomenclature of Territorial Units for Statistics) 3 level, corresponding to small regions for specific diagnoses, and were already provided as monthly aggregated data across years. We filtered the data for the years 2008 to 2019, with 2008 marking the start of the infection reporting period and 2019 selected to align with the most recent available observed environmental data. For further analysis, infection records were converted from NUTS3-level case data into spatially explicit observations on a 0.5° grid using centroid-based point locations for each affected NUTS3 municipality. Records potentially introducing spatial uncertainty (i.e., stemming from NUTS3 municipalities spanning more than one 0.5° cell) were excluded, and duplicate records within the same grid cell and month of a given year were removed.

For TBEV and WNV occurrences, we generated absence data by sampling infection-free cells within NUTS3 municipalities across EU/EEA countries with mandatory surveillance systems, using a balanced ratio to the presences. To ensure consistency with our temporal resolution, the absences were temporally matched to virus occurrences at a monthly timescale. Finally, to reduce spatial autocorrelation, we applied a 50 km thinning threshold (Aiello-Lammens et al., 2015) to both the monthly infection presence data and the corresponding absence data. Supplementary Figure S1 shows a map of all presence points for the viruses and the generated absence points.

### Environmental data

Climate data and land-use data were obtained from ISIMIP3 (Inter-Sectoral Impact Model Intercomparison Project Phase 3; https://www.isimip.org/protocol/3/; Frieler et al., 2024). From ISIMIP3a, we used historical simulations and counterfactual simulations that assume either no climate change or no land-use change (baseline year 1901), enabling the detection and attribution of observed impacts (Lange et al., 2025; Volkholz & Ostberg, 2024), while future projections were retrieved from ISIMIP3b (Lange & Büchner, 2021). For climate data, we focused on five key variables: mean near-surface air temperature (tas), minimum near-surface air temperature (tasmin), maximum near-surface air temperature (tasmax), precipitation (pr), and near-surface relative humidity (hurs). The data were provided at 0.5° spatial resolution, and daily outputs were aggregated into monthly values to match the temporal resolution of the analyses. Historical climate data included simulations reflecting observed climate change as well as counterfactual climate simulations based on detrended data from the GSWP3-W5E5 model (Frieler et al., 2024). Future projections were obtained under three different climate model-socioeconomic pathway combinations (SSP1-RCP2.6, SSP3-RCP7.0 and SSP5-RCP8.5) from the five available global climate models. For land-use data, we used historical datasets from ISIMIP3a, which stem from the LUH2 project (Land Use Harmonization project; version 2; Hurtt et al., 2020), comprising observed land-use reconstructions and a counterfactual scenario assuming no land-use change at annual time steps and 0.5° spatial resolution. Future land-use projections corresponding to the three SSP scenarios were sourced directly from LUH2, as ISIMIP3b did not yet include projected land-use states. Land-use categories were grouped into eight distinct classes for each year: primary forest, primary non-forest, secondary forest, secondary non-forest, pastures, rangelands, croplands, and urban areas. Future land-use data were aggregated to the 0.5° target spatial resolution (originally 0.25°). All environmental data were processed for the historical period from 1970 to 2019 and for the future period from 2030 to 2059. Note that the historical period extends beyond the temporal coverage of the available infection data. We have chosen this longer period to compare how predicted disease spread and phenology have changed over several decades and are expected to continue changing in the future. The associated implications for projected climate and land-use change across Europe under the three scenarios are summarised in Supplementary Table S2.

For the final input dataset used in the spatiotemporal SDMs for the main vector species, we matched species data, including presence observations and background data, with the month- and year-specific climate predictors, as well as year-specific land-use predictors. For the spatiotemporal virus models, infection data comprising reported presences and derived absences were matched exclusively with month- and year-specific climate predictors, as land use was not assumed to have any direct effect on virus prevalence. Details on the final species and infection occurrence numbers included in the models are provided in the Supplementary Table S3.

### Spatiotemporal species distribution models

We built nested, spatiotemporal SDMs using an ensemble approach. For the vector species, we included month-specific climate data and year-specific land-use data as predictor variables, and we predicted monthly habitat suitability. Since the distribution of the studied pathogens depends both on climate and on the presence of their vector species, we included month-specific climate data as predictors in the virus SDMs, along with the predicted monthly vector habitat suitability (hereafter referred to as “vector suitability”), following a nested modelling approach (Chen et al., 2024; Rochat et al., 2020). Before building the models, we performed a variable selection by identifying highly correlated variable pairs (Spearman correlations |r|> 0.7) and only retaining the more important variable from each pair (Dormann et al., 2013). Variable importance was evaluated using explained deviance from univariate generalised linear models (GLM) and a 5-fold cross-validation approach (Zurell, Zimmermann, et al., 2020). Given that the three temperature variables (tas, tasmin, tasmax) are highly correlated but may differentially influence species distribution across seasonal conditions, we opted for averaging their impact rather than selecting a single temperature variable from the correlation-based variable selection process. Consequently, we created three distinct predictor sets, each containing the identified most important and weakly correlated predictors, along with one of the temperature variables.

For both the target vector species and target viruses, spatiotemporal SDMs were then fitted using four statistical algorithms, each applied to all three temperature-variant predictor sets: GLM, generalised additive models (GAM), random forest (RF), and boosted regression trees (BRT). For vector models, regression methods (GLM, GAM) used all background data with equal weighting, while machine-learning approaches (RF, BRT) were trained on replicated balanced datasets (Barbet-Massin et al., 2012). Virus models were fitted using the balanced 1:1 presence-absence ratio without predefined weighting. Ensemble predictions were generated by averaging outcomes across predictor sets and algorithms, with additional averaging across replicates for vector machine-learning models. The performance of all spatiotemporal SDMs was assessed using a 5-fold cross-validation, with the area under the receiver operating characteristic curve (AUC) and the Boyce index (Hirzel et al., 2006) as focal evaluation metrics. We regard models obtaining an AUC or Boyce index above 0.7 as exhibiting fair predictive performance (Guisan et al., 2017). Partial response curves derived from cross-validated ensemble predictions for the vectors and their associated viruses are displayed in Supplementary Figure S2.

For future projections, ensemble predictions were averaged across the five different climate models under the same socio-economic scenario, thereby accounting for uncertainty among climate projections. In our manuscript, we present results exclusively for the SSP3-7.0 scenario, as it is considered to most realistically reflect current global policy trajectories (IPCC, 2021). Results based on the SSP1-2.6 and the SSP5-8.5 scenarios are provided in the Supplementary Material. Binary ensemble predictions were generated by applying the corresponding maxTSS (Maximising the True Skill Statistic) threshold proposed by Liu et al. (2013). All analysis outputs considered the study area encompassing EU/EEA countries (as of 2019), where surveillance data were largely available and reported to the ECDC, ensuring reliable results that are directly actionable within the coordinated EU/EEA public health system. Although predictions of the main vector species were not spatially restricted by administrative boundaries, the same study area was applied consistently to ensure comparability across results. To further enhance the ecological realism of our virus predictions, we masked (i.e., set to 0) all grid cells in the corresponding monthly predictions where the respective vector species was predicted to be absent, thereby ensuring that disease transmission risk was only considered in areas where transmission was biologically plausible. For WNV, the analysis was restricted to the months May through December, as historical infection data were only available for these months.

### Phenological analyses of vector activity and disease transmission risk

All phenological analyses were based on monthly habitat suitability ensemble predictions for *Ixodes ricinus*, TBEV, *Culex pipiens*, and WNV, covering both historical (1970 – 2019) and future (2030 – 2059) time periods. The historical period was selected to enable the long-term reconstruction of past suitability patterns, whereas the future period represents a policy-relevant mid-century projection window.

We assessed decadal trends in the timing of vector and virus suitability throughout the year, clearly separating a historical attribution step and a future projection step. In the attribution step, we aimed to disentangle the relative impacts of climate and land-use changes by comparing decadal phenology trends under factual (observed) and counterfactual climate and land-use simulations. For this, we compared ensemble predictions reflecting a mix of observed environmental conditions and counterfactual simulations: 1) Factual historical climate and land-use changes, capturing the realised environmental dynamics over time, 2) a historical counterfactual simulation assuming no climate change throughout the 20^th^ century, isolating the influence of climate, 3) a historical counterfactual simulation assuming no land-use change throughout the 20^th^ century, isolating the influence of land use, 4) a combined historical counterfactual simulation, in which both climate and land-use conditions remained fixed at pre-industrial 1901 values (Frieler et al., 2024). In the projection step, we projected potential future distributions of vectors and viruses under three future climate and land-use scenarios, representing different potential environmental trajectories. While land-use variables were not directly included in the virus models, the effects of land-use change were implicitly accounted for through the incorporation of the corresponding habitat suitability predictions of the main vector species. Results from the attribution and projection step were analysed at the European scale and at finer regional scales. First, for each target species and associated virus, we extracted monthly mean and peak occurrence probabilities per decade from the ensemble predictions, focusing specifically on the historical decades 1970s, 1990s, and 2010s, and the future decades 2030s and 2050s. To do so, we determined the monthly mean and peak suitability predictions across the entire study area for each year, where peak occurrence was defined as the 95^th^ percentile of predicted suitability values. Subsequently, we averaged these monthly values across all years within a given decade.

Second, we investigated how decadal trends in the seasonal timing of vector and virus suitability vary across Europe’s main climatic zones. We separately predicted decadal phenology trends for seven Köppen-Geiger climate zones that capture the major climate in Europe: Arid, Temperate (Mediterranean, Humid subtropical, Oceanic), Continental (Humid, Subarctic), and Polar. As climate and thus climate zones have considerably shifted in Europe over the past decades, we used for each target decade the corresponding Köppen-Geiger classification map at a spatial resolution of 0.5° (https://www.gloh2o.org/koppen; Beck et al., 2023): the 1961 – 1990 map for the 1970s, the 1991 – 2020 map for the 2010s, and the 2041 – 2070 maps for the 2050s, under the three projected socio-economic scenarios. We excluded climate classifications that were consistently represented by fewer than 15 cells across Europe. This approach allowed us a more nuanced assessment of regional and climate-specific phenology patterns across Europe’s climatic gradient and their development over time.

Lastly, we assessed the temporal trends in the duration of vector activity and potential disease transmission periods over the year and how these patterns may be changing across time and space. As in the previous analysis, we focused on the ensemble predictions based on factual historical climate and land-use changes, as well as on the three future socio-economic scenarios. This analysis was conducted using binarised predictions, indicating predicted presence or absence for each month. For each prediction year, we calculated the number of months with predicted presences per 0.5° cell, which we defined as the annual activity or transmission duration for that location. Based on these yearly values, we then computed the mean duration per decade for each cell. To assess future trends, we fitted a linear model for each cell using the decadal mean durations from the 2010s to the 2050s, based on the three future socio-economic scenarios. The slope of these models served as an indicator of change, allowing us to evaluate whether the activity or transmission duration of vectors and associated viruses is projected to increase, decrease, or remain stable in the coming decades.

For all analyses and visualisations, R 4.3.1 was used, along with the following packages: *CoordinateCleaner* (Zizka et al., 2019)*, dismo* (Hijmans et al., 2023)*, gbm* (Greenwell et al., 2022)*, mgcv* (Wood, 2003)*, PresenceAbsence* (Freeman & Moisen, 2008)*, randomForest* (Liaw & Wiener, 2002), and *terra* (Hijmans, 2023).

## Results

### Performance of spatiotemporal SDMs

Ensemble models of *Ixodes ricinus*, TBEV, *Culex pipiens*, and WNV demonstrated fair to good overall predictive performance, with AUC values of 0.73, 0.76, 0.74, and 0.83, respectively. This finding was further supported by the consistently high Boyce index values, close to or equal to 1. Monthly model performance showed greater variability. For *Ixodes ricinus*, AUC values ranged from 0.59 in January to 0.79 in March, while for TBEV, performance ranged from 0.57 in March to a maximum of 0.85 in August. *Culex pipiens* predictions showed the lowest AUC in January (0.65) and the highest in February (0.82). Cross-validated predictions for WNV performed well throughout the studied months, with AUC values above 0.77. Nevertheless, for each of the four target species and pathogens, AUC values remained predominantly within a fair to good performance range across months, with values falling below the commonly used 0.7 threshold in no more than five months, and only rarely falling substantially below this level. Monthly Boyce index values also predominantly exceeded a value of 0.7. A more detailed performance overview is provided in Supplementary Table S4.

### Historical and future risk distribution patterns

We first assessed distributional shifts in TBEV and WNV transmission risk across Europe from the 1970s to the 2050s, focusing on the months of highest predicted transmission risk, May and July, respectively (Figure 1). For TBEV, transmission risk became increasingly spatially clustered from the 1970s to recent decades, with hotspot regions predicted to expand and intensify, particularly in central and eastern Europe, by the 2050s under the SSP3-7.0 scenario (Figure 1; left panels). The spatial risk distribution of virus transmission largely overlapped with the predicted distribution of *Ixodes ricinus*, suggesting that TBEV is primarily constrained by its vector. For WNV, transmission risk showed a clear expansion and increase from the 1970s to the 2010s across large parts of Europe, including central, eastern, southern, and northeastern regions (Figure 1; right panels). These hotspots are projected to further consolidate by the 2050s under the SSP3-7.0 scenario, with particularly high predicted WNV suitability in, for instance, northern Italy, and parts of Romania, Portugal, and Finland. In contrast, virus transmission risk is projected to remain consistently limited in northern and northwestern Europe, indicating that the potential distribution of WNV is more constrained than that of its vector species, *Culex pipiens*.

**Figure 1:**
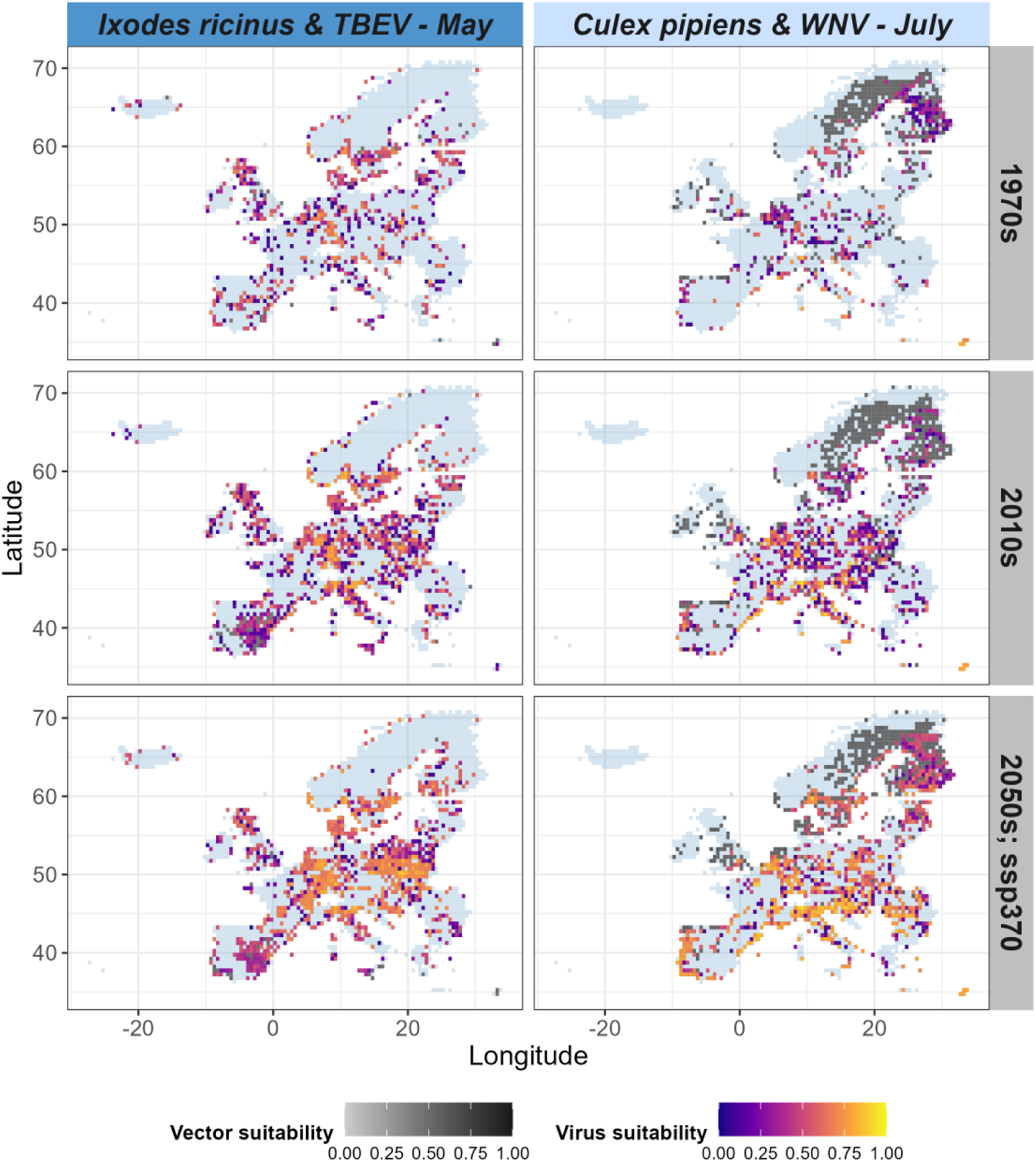
Predicted distributions of vector activity and virus transmission risk over time across Europe (EU/EEA countries with UK). Distribution trends of *Ixodes ricinus* and Tick-borne encephalitis virus (TBEV; left panels) are shown for May, while trends of *Culex pipiens* and West Nile virus (WNV; right panels) are shown for July. Risk distributions are illustrated for three target decades: 1970s, 2010s, and 2050s, with future projections based on the SSP3-7.0 scenario. Only grid cells with vector or virus suitability are displayed where the vector or the virus were predicted to be present within the selected month of a given decade. Predicted presences were determined using the threshold that maximises the True Skill Statistic (maxTSS). Predicted habitat suitability values for the vectors and viruses were averaged across each decade. Vector suitability is represented using a grey scale, with virus suitability overlaid using a coloured scale.

Under both alternative future scenarios, SSP1-2.6 and SSP5-8.5, TBEV and WNV hotspots are projected to expand markedly compared with the SSP3-7.0 scenario. In particular, regions in central and eastern Europe are expected to experience the strongest increase in transmission risk, forming large, spatially connected areas of high virus suitability (Supplementary Material, Figures S3+S4).

### Historical phenological patterns in risk intensity

Historical seasonal risk curves for *Ixodes ricinus* and its associated virus, TBEV, showed a marked increase in risk levels across Europe from the 1970s to the 2010s, accompanied by a clear shift in seasonal patterns (Figure 2; first and second rows). We found two major changes to characterise this transformation: first, the seasonal phenology shifted from a predominantly unimodal pattern with a broad, flat peak to a distinctly bimodal curve with two pronounced peaks, one in spring (May) and another in autumn (September). Second, the timing of the spring peak shifted to earlier in the year, from June to May. This shift was particularly notable for TBEV, especially between the 1990s and 2010s. Counterfactual model predictions indicated that these historical changes were strongly driven by climate and land use changes. When 20^th^-century anthropogenic influences were excluded, risk levels remained considerably lower, and the seasonal profile retained a simpler, predominantly unimodal curve. Among the drivers, climate change emerged as the main driver of the observed bimodal risk pattern. In the simulation with detrended climate, the characteristic mid-summer decline in mean occurrence probabilities did not occur, indicating that this seasonal dip is a consequence of climate-driven shifts. In contrast, land-use had impacted the overall risk levels with considerably lower mean suitability under the counterfactual land-use simulations.

**Figure 2:**
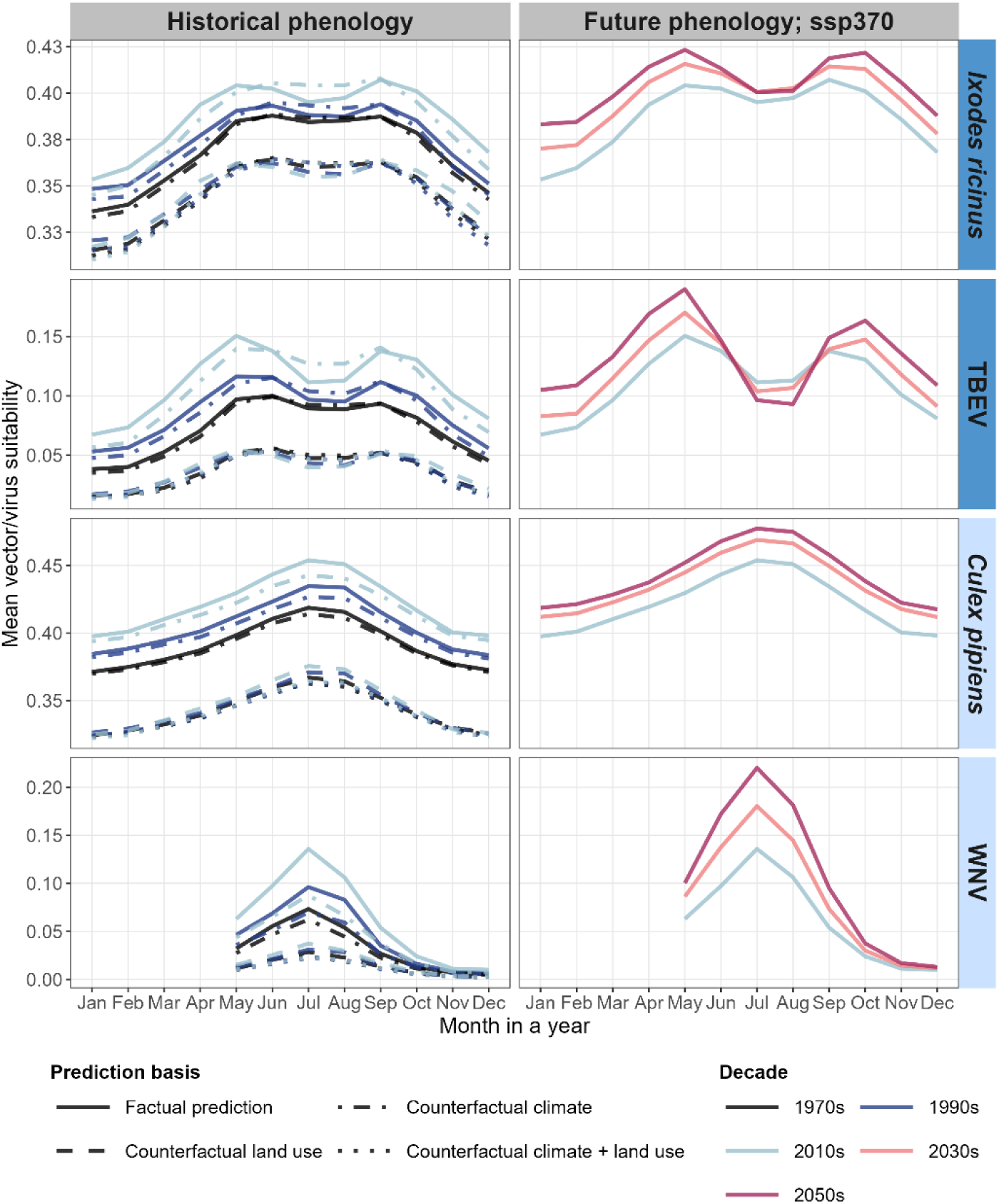
Predicted seasonal risk curves of vector activity and virus transmission across historical and future decades. Decadal trends in the seasonal phenology of activity risk for *Ixodes ricinus* and *Culex pipiens* are shown (top and third rows), along with the transmission risk for their associated viruses, Tick-borne encephalitis virus (TBEV) and West Nile irus (WNV) (second and fourth rows). Risk is represented as the average monthly predicted habitat suitability across Europe (EU/EEA countries with UK). Line colours distinguish decades (1970s, 1990s, 2010s, 2030s, 2050s), with phenologies of past decades shown in the left column and future decades in the right column. Line styles differentiate between factual and counterfactual environmental simulations (reference year 1901), including counterfactual climate, counterfactual land use, and their combination. Future projections are based on the SSP3-7.0 socio-economic scenario. For WNV, seasonal risk curves from January to April are not shown due to the absence of historical infection reports during these months.

For *Culex pipiens* and its associated virus, WNV, historical seasonal risk curves across Europe also showed a clear upward trend in both vector activity and transmission risk (Figure 2; third and fourth rows). Their phenology was characterised by a pronounced summer peak, with risk levels reaching their maximum in July. Counterfactual model simulations demonstrated that these historical increases were strongly attributable to climate and land-use changes. Compared to *Culex pipiens*, WNV displayed a considerably steeper phenological curve, with transmission risk declining more rapidly toward spring and autumn.

### Future phenological patterns in risk intensity

For *Ixodes ricinus* and its associated pathogen, TBEV, future projections under the SSP3-7.0 scenario suggest a continued rise in overall risk levels, with increasingly sharp seasonal fluctuations, especially for TBEV. Between the 2010s and the 2050s, the bimodal pattern is projected to intensify across Europe, with the spring peak becoming more dominant than the autumn peak. Moreover, both *Ixodes ricinus* and TBEV show a shift to later autumn peaks, from September to October. Interestingly, projections for the 2050s indicate a further decline in TBEV risk levels during the summer months, dropping even below those predicted for previous decades.

The pronounced summer peak in the suitability of *Culex pipiens* and its associated pathogen, WNV, is projected to intensify under future conditions. Specifically, under the SSP3-7.0 scenario, WNV suitability for the July peak is expected to increase from 0.14 in the 2010s to 0.22 in the 2050s. For both *Culex pipiens* and WNV, we found that rising future risk levels are closely linked to a widening of the phenology curve, indicating an extended seasonal window of activity and transmission risk across Europe in the coming decades.

Projected phenology curves under the SSP1-2.6 and SSP5-8.5 scenarios closely resemble those of the SSP3-7.0 scenario. Notably, under the SSP1-2.6 scenario, TBEV transmission risk is projected to be comparably more pronounced, with the mean suitability reaching 0.27 during the first seasonal peak in May by the 2050s (Supplementary Material, Figure S5). A similar intensification of TBEV risk is also evident under the SSP5-8.5 scenario. Furthermore, SSP5-8.5 projections suggest a more substantial increase in WNV transmission risk compared to SSP3-7.0, with the mean suitability reaching 0.29 at the July peak by the 2050s (Supplementary Material, Figure S6). When examining peak suitability as an indicator of vector activity and disease transmission risk, we found that decadal risk levels for both TBEV and WNV are projected to reach maximum decadal values of 0.73 and 0.79, respectively, by the 2050s. These values markedly exceed the corresponding decadal means (0.19 for TBE, 0.22 for WNV), hinting at substantial spatial heterogeneity of transmission risk with localised hotspot areas across Europe (Supplementary Material, Figures S7–S9).

### Historical and future phenological patterns in risk intensity across space

Model predictions for *Ixodes ricinus* suggested that Temperate and Arid climate zones exhibited consistently high activity risk throughout the year during the past decades of the 1970s and the 2010s (Figure 3; second row). However, within Mediterranean and Humid Subtropical regions, a pronounced decline in activity risk was predicted during the summer months, marking their lowest seasonal risk within the year. While future projections across these climate zones indicate an overall increase in mean suitability by the 2050s, this characteristic summer decline is expected to become increasingly pronounced and prolonged. In contrast, the other climate regions were found to follow a unimodal phenological pattern. Among these, predictions for Continental Humid climates, spanning much of Central and Eastern Europe, were associated with comparably high vector activity risk, especially during the peak months from May to September in the 2010s, when suitability for *Ixodes ricinus* reached some of the highest levels predicted across Europe. Looking ahead to the 2050s, no substantial increase in activity risk is projected for these regions. This may be due to the projected northward shift and spatial contraction of continental subhumid zones in Eastern Europe (Figure 3; first row). Polar and Continental Subarctic regions, in particular, maintained comparably lower seasonal occurrence probabilities since the 1970s, with no substantial increases anticipated under the SSP3-7.0 climate scenario.

**Figure 3:**
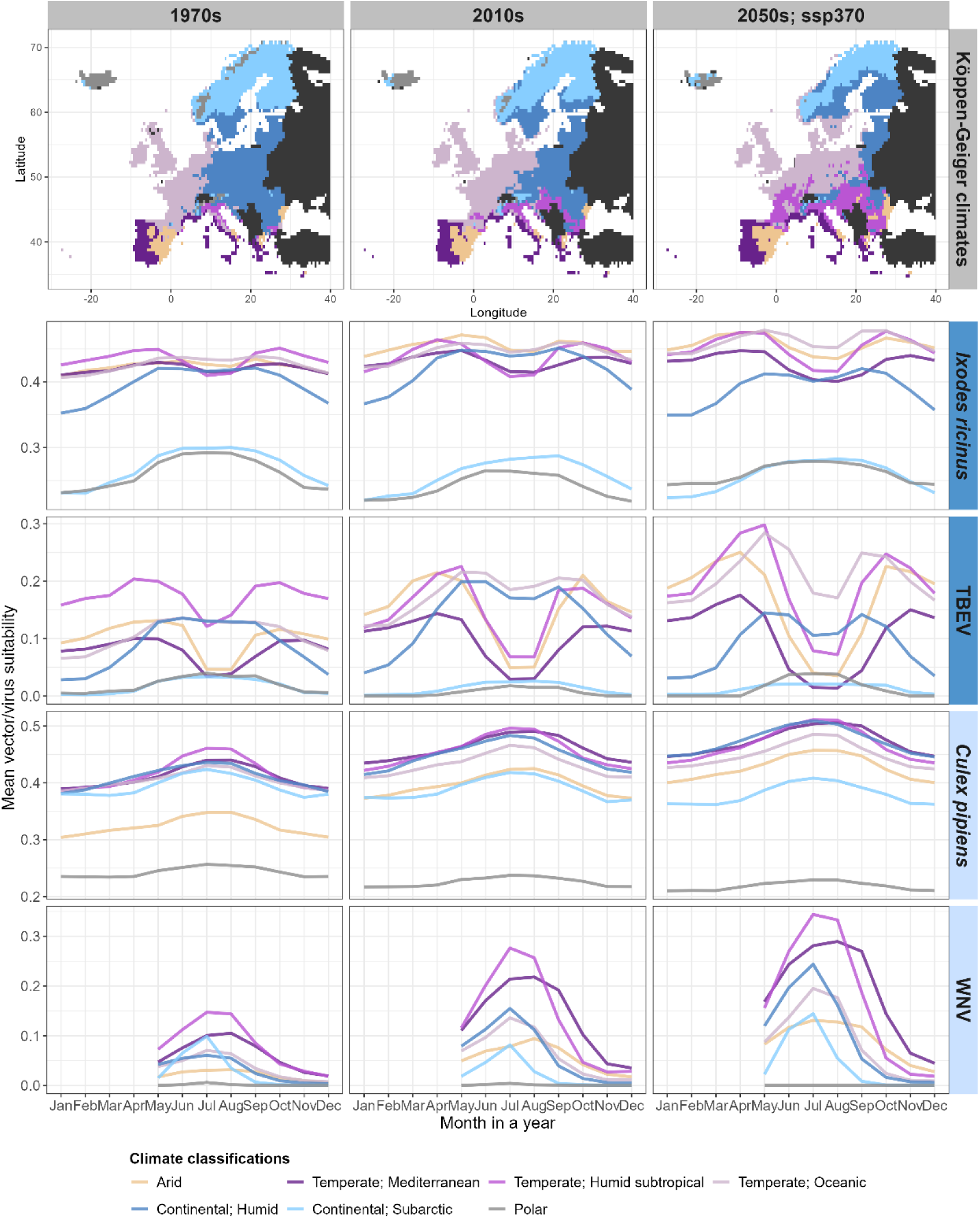
Predicted seasonal risk curves of vector activity and virus transmission by Köppen-Geiger climate regions over time. The top row shows the spatial distribution of seven different Köppen-Geiger classifications across Europe (EU/EEA countries with UK) corresponding to three time periods that align with the decades studied: 1961-1990 (representing the 1970s), 1991 – 2020 (2010s), and 2041 – 2070 (projected 2050s under the SSP3-7.0 scenario). The rows below display the seasonal phenology of vector activity risk for *Ixodes ricinus* and *Culex pipiens* (second and fourth rows), along with the transmission risk of their respective viruses, Tick-borne encephalitis virus (TBEV) and West Nile virus (WNV) (third and fifth rows). Vector activity and transmission probability are expressed as the average monthly predicted habitat suitability across the prevailing Köppen-Geiger climate zones in each respective time period. Line colours indicate the different Köppen-Geiger classifications: Arid, Temperate (Mediterranean, Humid subtropical, Oceanic), Continental (Humid, Subarctic), and Polar. For WNV, seasonal risk curves from January to April are not shown due to the absence of historical infection reports during these months.

For TBEV and its primary vector, *Ixodes ricinus*, the highest transmission risks in the past decades were predicted for Arid and Temperate climate zones, particularly during the early spring and late autumn months, with a notable decline during the summer (Figure 3). An exception to this pattern was found in Temperate Oceanic climates, where predictions followed a rather unimodal seasonal profile. Especially during the 2010s, retrospective predictions suggested that both Temperate Oceanic and Continental Humid regions sustained the highest TBEV transmission risks during the summer months, from May to September. By the 2050s, the most substantial increases in transmission risks are projected for the Temperate Humid Subtropical, Temperate Oceanic, and Arid climate regions. These areas are expected to experience a lengthened transmission season, with elevated risk beginning earlier in spring and extending further into autumn. However, the characteristic summer decline in risk is also projected to become more prolonged. Notably, in Temperate Oceanic climates, transmission risk is projected to remain comparably high throughout the year, with these regions shifting markedly northeastward under the SSP3-7.0 future scenario (Figure 3; first row). As with its primary vector, *Ixodes ricinus*, transmission risk in Continental Subarctic and Polar climate zones was predicted to remain consistently low, with no substantial increases anticipated.

*Culex pipiens* predictions showed a consistent increase in activity risk from the 1970s to the 2050s across nearly all climate zones (Figure 3; fourth row). By the 2050s, the highest activity risks throughout the year are projected for Continental Humid and the three Temperate climate types. As climate zones continue to shift across Europe (Figure 3; first row), the potential seasonal range of *Culex pipiens* is projected to expand northward. In general, the increasing activity risk is accompanied by a broadening of the seasonal activity window, suggesting an extended period of mosquito activity in the future decades compared to historical patterns.

Patterns of WNV transmission risk revealed a clear increase from the 1970s to the 2050s across all climate zones, except for Polar regions (Figure 3; fifth row). We found these increases to be especially pronounced in Temperate and Continental climate zones. Among these, regions classified as Temperate Humid Subtropical exhibit the highest overall WNV transmission risk values in all decades. In contrast, Temperate Mediterranean climates showed the most prolonged period of potential high WNV transmission under current and future environmental conditions, characterised by the broadest phenology curve. Interestingly, in these Mediterranean regions, the peak transmission risk is projected to occur in August, one month later than in most other climate zones.

Under both alternative future scenarios, the Temperate Oceanic climate regions are projected to sustain even higher risk levels for TBEV and WNV, and their respective primary vector species, *Ixodes ricinus* and *Culex pipiens* (Supplementary Material, Figures S10+S11). Furthermore, phenology curves using peak suitability as a risk indicator exhibit only minor deviations in magnitude from the mean suitability, suggesting that risk is more evenly distributed within the main climate regions across Europe (Supplementary Material, Figure S12-S14).

### Future phenological patterns in risk duration

Under the SSP3-7.0 scenario, we found that hotspots of prolonged *Ixodes ricinus* activity are projected to emerge across various regions of Europe (Figure 4; first row). These include parts of Eastern Europe, particularly Poland and the Czech Republic, as well as areas in Northern Europe, such as southern Finland and southern Sweden. Additionally, southwestern Europe is expected to be affected, including regions of Spain and the border zone between France and Germany. Within these emerging hotspots, the tick activity season is projected to lengthen by up to 0.36 months per decade, equivalent to an expansion of roughly 11 days every ten years, continuing through to the 2050s. TBEV exhibited an almost identical pattern of future spatial hotspots (Figure 4; second row), with 31.3 % of the EU/EEA area projected to experience a prolonged transmission period, compared to 32.5 % for *Ixodes ricinus*. However, finer regional differences became evident when comparing the retrospectively predicted length of the TBEV transmission season and the *Ixodes ricinus* activity season in the 2010s. Notably, in parts of southern Europe, including Spain and Italy, as well as the northern UK, the transmission season of TBEV remained shorter than the activity season of its main vector, *Ixodes ricinus*. This difference is further reflected in the average duration of risk periods during the 2010s, with *Ixodes ricinus* showing a mean activity season of

**Figure 4:**
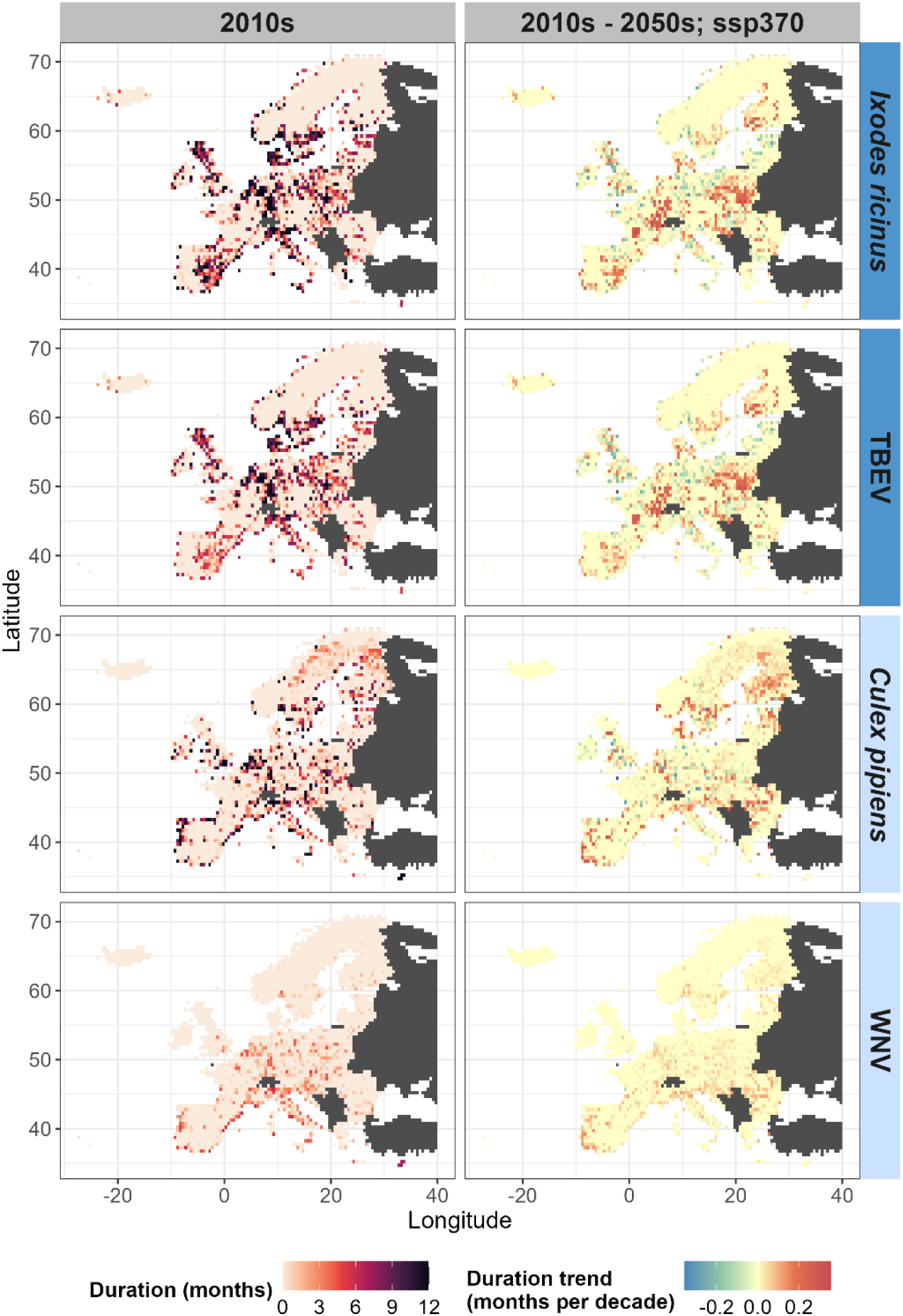
Predicted duration and temporal trends of seasonal risk periods for vector activity and virus transmission across Europe (EU/EEA countries with UK). The left column maps the average duration (in months) of seasonal risk during the 2010s for *Ixodes ricinus* and *Culex pipiens* (first and third rows), and their associated viruses, Tick-borne encephalitis virus (TBEV) and West Nile virus (WNV) (second and fourth rows). Darker shades in the left column indicate longer risk periods, up to 12 months. The right column shows the respective projected trends in seasonal risk duration for each species and virus presented in the left column, from the 2010s to the 2050s under the SSP3-7.0 scenario, expressed as the rate of change in months per decade. Warmer (yellow-red) and cooler (blue-green) tones in the right column represent projected increases or decreases in risk duration, respectively. Seasonal risk durations are derived from binary maps of monthly predicted habitat suitability, thresholded to maximise the True Skill Statistic (maxTSS).

### 2.3 months across Europe, compared to 1.9 months for TBEV

For *Culex pipiens*, a prolonged activity period under the SSP3-7.0 scenario is expected across large parts of Europe, particularly in northern regions including Sweden, Finland, Estonia, and Denmark (Figure 4; third row). In these areas, the activity season is projected to lengthen by up to 0.35 months per decade, corresponding to an approximate increase of 10.7 days every ten years. Similar patterns are anticipated in parts of southern and southeastern Europe, for example, in northern Italy and its eastern neighbours, including Slovakia, Hungary, and Romania, as well as areas of the Iberian Peninsula, including parts of Spain and Portugal. In contrast, both the past and projected future ranges of WNV appeared more limited than that of its primary vector, *Culex pipiens* (Figure 4; fourth row). In northern Europe, WNV seemingly approached the limits of its potential transmission range, with comparably little to no seasonal presence predicted in countries such as Sweden, Denmark, the UK, and Ireland. This distinction became evident in predictions for the 2010s, which estimated that 71.0 % of the EU/EEA area had no seasonal WNV transmission risk, compared to 54.4 % for *Culex pipiens.* Nevertheless, a general lengthening of the WNV transmission season is anticipated in many parts of Europe, largely reflecting the same regions where extended vector activity is predicted for *Culex pipiens*. Specifically, a future lengthening of the transmission period is predicted for 31.7 % of the EU/EEA area, compared to 40.7 % for *Culex pipiens*.

Projected hotspots of prolonged activity under the SSP1-2.6 and SSP5-8.5 scenarios show notable deviations compared to those identified under SSP3-7.0 (Supplementary Material, Figures S15+S16). Importantly, under both alternative scenarios, central Europe, particularly Germany, is projected to emerge as a key transmission hotspot for both TBEV and WNV. Additionally, northern Europe, especially regions of Sweden and Finland, is projected to be more strongly affected by a lengthening of the seasonal window of activity and transmission for *Ixodes ricinus*, TBEV, *Culex pipiens*, and WNV. This trend is further reflected in the projected increase in the proportion of area expected to undergo an extended risk period across the EU/EEA area. When comparing SSP3-7.0 with the maximum values observed under SSP1-2.6 and SSP5-8.5, we found the largest increases in area affected were +14.7 % for *Ixodes ricinus*, +14.2 % for TBE, +11.6 % for *Culex pipiens*, and +2.3 % for WNV.

## Discussion

Within this study, we examined how the spread and phenology of TBEV and WNV respond to changing climate and land-use conditions across Europe, providing a basis for anticipating spatiotemporal trends in infection risk and thereby guiding effective disease risk management. Historical attribution analyses indicate that land-use change primarily shaped the spatial distribution of the vectors and thus transmission risk, whereas climate change was the dominant driver of seasonal dynamics and phenological shifts. Our findings highlight a clear future trend: both diseases are projected to impose an increasing burden over the coming decades. Europe-wide transmission risk is expected to rise for both TBEV and WNV, and TBEV is additionally projected to undergo pronounced phenological shifts, characterised by a dominant spring peak and a delayed autumn peak, extending into October. This elevated transmission risk is further associated with prolonged seasonal transmission windows. Importantly, these hotspots of extended transmission are projected to intensify as well as to expand spatially across large regions of Europe. We discuss these findings in light of public health relevance and intervention planning to effectively manage the anticipated increase in TBEV and WNV burden across Europe.

We implemented a robust modelling framework to understand the spatiotemporal dynamics of TBEV and WNV under changing climate and land-use conditions over four decades of historical changes and into the future. Our nested SDM approach jointly considered vectors and viruses and, combined with a detection and attribution framework (Gonzalez et al., 2023; Schrodt et al., 2025; Thomas et al., 2026), proved to be a powerful tool for disentangling the relative contributions of environmental drivers to past patterns of transmission risk. Our results indicate that historical land-use change primarily contributed to increases in absolute vector suitability, potentially shaping where transmission is most likely to occur, whereas phenological dynamics were largely driven by climatic factors, suggesting that climate determines when transmission is possible and for how long. This distinction became particularly evident in the predicted summer dip in TBEV transmission risk, which was primarily attributable to climatic changes. Importantly, these strong absolute and direct effects of land-use change on vector suitability may be mediated by intermediate hosts: for example, rodents, which serve as key reservoirs for TBEV, could benefit from land-use changes such as afforestation and reforestation, as well as crop-to-grass conversions associated with land abandonment, processes that dominated land-use change over the last century (Fuchs et al., 2015). Such habitat modifications may have increased rodent abundance and distribution, subsequently enhancing the population of *Ixodes ricinus* and thereby amplifying the risk of TBEV transmission (Dagostin et al., 2023; Friedsam et al., 2022). Similarly, for WNV, historical land-use changes, including urbanisation and cropland expansion (Fuchs et al., 2015), may have created habitats favourable to both synanthropic passerine birds, which are considered key amplifying hosts in Europe (Simonin, 2024), and competent mosquito vectors such as *Culex pipiens*. This overlap in human-modified landscapes may have facilitated viral amplification and WNV spillover into human populations in the past (Giesen et al., 2023; Kilpatrick, 2011; Marcantonio et al., 2015).

Historical predictions for TBEV and WNV, along with their respective main vector species, revealed stark differences in the phenological dynamics of these diseases across Europe. For TBEV, our historical predictions indicated that *Ixodes ricinus* activity and the corresponding transmission risk shifted from a unimodal to a bimodal curve, with two distinct seasonal peaks occurring in spring and autumn. Previous studies have demonstrated that this characteristic pattern is driven by temperature as a primary trigger of tick activity, while the saturation deficit, a measure of air dryness combining temperature and relative humidity, acts as a key within-season regulator (Schulz et al., 2014; Zając et al., 2021). Saturation deficit typically reaches its maximum during mid-summer (July-August) in Europe, coinciding with periods of reduced questing activity of *Ixodes ricinus* due to elevated desiccation risk (Schulz et al., 2014; Zając et al., 2021). Similarly, Lamsal et al. (2023) reported that TBEV prevalence in ticks is negatively affected by lower relative humidity, likely further contributing to the reduced transmission risk when tick activity is already constrained. Notably, while our historical predictions suggested that the spring and autumn peaks in *Ixodes ricinus* activity risk were of comparable magnitude, TBEV transmission risk was predicted to have developed a progressively stronger spring peak over time. For *Ixodes ricinus*, these predictions are only partially consistent with historical field observations across Europe, which generally report the highest tick abundances in late spring or early summer, exceeding those of the secondary autumn peak (Bartosik et al., 2011; Borde et al., 2019). This observed peak asymmetry in tick abundance has further been associated to seasonal increases in rodent host populations, which enhance host-tick encounter rates and subsequent feeding success (Cayol et al., 2017). As our modelling framework did not explicitly account for host-tick interactions, such biotic drivers may explain the described discrepancies between historically predicted activity risk in our study and the field-observed tick abundance patterns. Nevertheless, the predicted peak asymmetry between TBEV transmission risk and vector activity risk highlights that transmission is not solely determined by tick activity, but is strongly sensitive to climatic conditions, in line with recent studies (Daniel et al., 2018; Lamsal et al., 2023). Reflecting this climatic influence, our historical risk predictions of TBEV transmission in particular indicate that both the timing and relative magnitude of seasonal peaks vary across Europe’s major climate zones. In southern regions – characterised by Arid, Temperate Mediterranean, and Temperate Humid Subtropical climates – our retrospective predictions showed a pronounced bimodal seasonal pattern, with markedly reduced transmission risk during the summer months, consistent with observations from endemic areas in Croatia (Vilibic-Cavlek et al., 2024). In central and northern zones, including Temperate Oceanic and Continental Humid regions, historical predictions indicated persistently elevated transmission risk throughout the summer and a predominantly unimodal seasonal profile. Previous studies conducted within these climate zones have reported heterogeneous phenological patterns, even at small spatial scales. While TBEV infections have been observed as exhibiting a strong unimodal peak in June-July (Hellenbrand et al., 2019; Kriz et al., 2012), *Ixodes ricinus* occurrence has been described to vary between unimodal and bimodal patterns. These differing patterns have been attributed to variation in tick developmental stage and habitat-specific microclimatic conditions, such as near-ground temperature (Borde et al., 2021; Daniel et al., 2015; Wilhelmsson et al., 2013). Taken together, despite fine-scale ecological variability observed in localised field studies, our historical predictions indicate that large-scale climate has been the dominant driver of continental TBEV seasonality, implying the potential to time and target public health measures according to regional climate-driven risk periods.

For WNV, our Europe-wide historical predictions displayed a distinctly different seasonal profile: *Culex pipiens* activity risk and the corresponding transmission risk were characterised by a pronounced summer peak in July, reflecting the role of higher temperatures in increasing mosquito population abundance, accelerating viral replication within the mosquito vector, and thereby enhancing virus transmission rates (Camp & Nowotny, 2020; Paz, 2015). The steeper seasonal profile of predicted WNV transmission risk relative to *Culex pipiens* activity risk is consistent with the strong temperature dependence of vector competence, which encompasses the mosquito’s overall ability to acquire, replicate, and transmit the virus. As demonstrated by Kilpatrick et al. (2008), WNV transmission rises sharply with temperature, constraining viral replication and effective transmission to a relatively narrow mid-summer period, even when mosquito activity persists. Interestingly, while WNV infections across Europe have generally peaked in August over the last decade (Camp & Nowotny, 2020; European Centre for Disease Prevention and Control, 2021), our historical risk predictions indicate that Europe-wide WNV transmission risk was highest in July. Importantly, we predicted the historical potential distribution of both the virus and its main vector, including areas predicted to be ecologically suitable where no infections have been reported, which may help explain discrepancies between predicted peak transmission risk and observed case timing. This possible explanation is reinforced by regional variation in seasonal patterns across major climate zones: in southern European regions with Temperate Mediterranean and Arid climates, such as Italy, Greece, Romania, and Croatia, where most WNV infections have been documented in the last decade (European Centre for Disease Prevention and Control, 2021), predicted transmission risk peaks in August rather than July. A further southern European climate zone, the Temperate Humid Subtropical region, exhibited a July peak, although predicted transmission risk remained elevated into August rather than declining sharply. In contrast to TBEV, historical predictions of WNV transmission risk exhibited a broadly consistent seasonal timing across climate zones, with summer peaks occurring in July or August, while climate primarily influenced the magnitude of risk. Accordingly, continental-scale WNV public health measures show the potential to be largely implemented within a shared seasonal window across regions, while adapting the scope of interventions according to climate-driven differences in transmission risk intensity.

Our future projections suggest that ongoing climate and land-use changes are likely to substantially intensify transmission risk for both TBEV and WNV across Europe. For TBEV, this intensification is projected to alter the seasonal profile of transmission risk, with the spring peak becoming increasingly dominant and the autumn peak shifting in timing, extending into October. Both TBEV and WNV are also projected to experience a longer period of elevated transmission risk. Extended transmission seasons, combined with elevated risk, not only increase the cumulative exposure but also amplify the likelihood of infection at any given time due to increased biting frequency of tick and mosquito vectors (Kilpatrick, 2011; Semenza & Paz, 2021). These projected changes render current surveillance systems, which are largely passive and unevenly coordinated across Europe, insufficient for effective disease control, particularly noting that both diseases can lead to serious neurological complications or even fatal outcomes (European Centre for Disease Prevention and Control, 2021; Heuverswyn et al., 2023; Wagner et al., 2020). Current WNV surveillance in EU/EEA countries relies predominantly on passive reporting of human infections, with weekly updates to ECDC during the transmission season and routine annual reporting. Animal health surveillance is implemented in 22 of the 30 countries, but is also largely passive, and integrated One Health approaches are implemented in only a minority of countries (European Centre for Disease Prevention and Control & European Food Safety Authority, 2023). In contrast, current TBEV surveillance focuses primarily on human cases and is largely based on annual passive reporting (European Centre for Disease Prevention and Control, 2022). Based on our findings, we suggest earlier initiation and prolonged continuation of weekly WNV surveillance reporting. Specifically, the current June-November window for weekly reporting (European Centre for Disease Prevention and Control & European Food Safety Authority, 2023) is too narrow and should be extended to ensure that surveillance covers the broader period during which transmission risk is expected to be elevated. Ideally, this approach should be complemented by integrated One Health surveillance, including active infection monitoring in vectors and non-human vertebrate hosts starting in spring, which can provide early warning of risk to humans weeks in advance (U.S. Centers for Disease Control and Prevention, 2024). Surveillance and early warning should be complemented by strengthened prevention and control interventions, including public awareness campaigns promoting individual prevention measures, both sustainable preventive and reactive vector control, and healthcare preparedness, including measures such as systematic screening of blood donations to prevent transfusion-associated WNV infections (European Centre for Disease Prevention and Control & European Food Safety Authority, 2023). For TBEV, the predicted shifts in seasonality may challenge prevention strategies that are based on historical exposure patterns. This aspect is particularly important because TBEV transmission is vaccine-preventable, underscoring the need to reconsider vaccination schedules and ensure completion of primary and booster doses before the start of the transmission season. Timely vaccination is especially critical, both because most TBEV infections occur in unvaccinated or incompletely vaccinated individuals (European Centre for Disease Prevention and Control, 2024) and because the spring transmission peak is projected to become increasingly dominant. Coordinating risk communication with these shifting seasonal risk patterns can further support timely vaccination uptake and reinforce preventive behaviour when exposure risk is highest.

Beyond the projected temporal shifts in phenology and increases in peak virus suitability, our results indicated the emergence of hotspots characterised by prolonged seasonal risk and a continued spatial expansion of TBEV and WNV through the 2050s. Several European regions are, therefore, likely to experience unprecedented risk levels in the future. Regions with increased WNV risks include Slovenia, Portugal, and areas of southern Finland and southern Sweden, underscoring the potential for further northward expansion of transmission risk. For TBEV, the further intensification and spread of hotspots within current endemic areas could strongly affect Poland, as well as southern Sweden, southern Finland, and areas along the France-Germany border. Localised areas within the Iberian Peninsula and the mainland area of Denmark, which have so far been only lightly affected, may emerge as new TBEV hotspots. Under both alternative socio-environmental futures, Central Europe, particularly Germany, which already has established TBEV endemic areas but has been only lightly affected by WNV, may experience a substantial increase in the burden of both diseases. Overall, these patterns highlight the need for context-specific preparedness, taking into account epidemiological history to distinguish between managing risk in endemic regions and the early detection of emerging transmission in previously low-risk areas. Our results can guide the implementation of monitoring and vaccination strategies in both endemic and previously unaffected regions.

### Limitations of the approach

Although providing important insights into the potential spread and phenology of TBEV and WNV under climate and land-use change, our results need to be interpreted cautiously for several reasons. First, the vector occurrence and disease data used are subject to inherent spatial and temporal biases (Bowler et al., 2022; European Centre for Disease Prevention and Control, 2019), potentially leading to uncertainties in the estimation of vector and virus niches over time. To reduce such spatial biases, we incorporated data filtering steps and spatial thinning to reduce the influence of clustered observations. Temporal bias related to seasonal extrapolation was addressed by restricting predictions to months for which observational data were available. Nevertheless, changes over time, such as improvements in diagnostic capacity and medical diagnostics, may have influenced reported incidence patterns, independent of actual transmission dynamics. For TBEV, incidence may further be influenced by vaccination coverage, as regions with higher vaccination rates are likely to show lower case numbers regardless of vector and virus suitability (Angulo et al., 2023). Second, the results of historical predictions should be viewed as showing potentially suitable areas and can, therefore, extend beyond known endemic areas. These deviations may arise from factors such as unaccounted biotic interactions and human intervention. In this regard, we only included the respective primary vector species for TBEV and WNV in Europe due to their dominant role in Europe-wide transmission. For WNV, while other mosquito species are also competent vectors, they occupy more limited geographic ranges, are only recently expanding, and contribute mainly to sylvatic rather than human transmission cycles (Camp & Nowotny, 2020). Likewise, for TBEV, other competent tick vectors, such as *Dermacentor reticulatus,* can contribute to transmission, but their impact is generally restricted to more localised areas (Ličková et al., 2020). Host species were also not explicitly included in our models. However, in Europe, the primary vectors feed on a wide range of hosts that are generally widespread and abundant (Chitimia et al., 2023; European Centre for Disease Prevention and Control & European Food Safety Authority, 2025), suggesting that host availability is rather unlikely to be a limiting factor at broader spatial scales. Third, the relatively coarse spatial resolution of 0.5° in our study may mask fine-scale suitable habitats, potentially overlooking areas of high TBEV or WNV transmission risk. Clearly, finer-resolution analyses could help identify these localised high-risk areas and provide a more accurate basis for targeted interventions.

## Conclusion

Future climate and land-use changes are projected to extend and intensify the seasonal risk of TBEV and WNV across Europe, with both endemic and previously unaffected regions likely to face increasing disease burden. Our projections suggest that WNV interventions could be coordinated continent-wide within a common, yet extended, seasonal window, while adapting the scope of interventions to regional transmission risk and epidemiological history. In contrast, TBEV interventions, in particular vaccination, may benefit from climate-informed scheduling of monitoring and mitigation measures aligned with shifting seasonal risk periods, again with region-specific adaptations. By identifying areas where transmission risk is expected to intensify or newly emerge, our results support the spatial prioritisation of surveillance, vaccination, and vector control strategies. More broadly, the continental perspective provided here underscores the importance of harmonised transnational cooperation for anticipating and managing climate-sensitive zoonotic diseases. Integrating spatiotemporal projections with coordinated public health planning can facilitate a shift towards upstream prevention that will be essential for the effective management of future disease burden of TBE and WNV in Europe.

## Author Contributions

Valén Holle and Damaris Zurell conceived the ideas and designed the methodology; Valén Holle prepared and analysed the data; Valén Holle led the writing of the manuscript; All authors contributed critically to the drafts and gave final approval for publication; Nadja Kabisch and Damaris Zurell secured the funding.

## Acknowledgements

We acknowledge funding support by the European Union through the project Zoonosis Emergence across Degraded and Restored Forest Ecosystems (project no. 101135094). This work further used resources of the Deutsches Klimarechenzentrum (DKRZ) granted by its Scientific Steering Committee (WLA) under project bb0820.

Disclaimer: The views and opinions of the authors expressed herein do not necessarily state or reflect those of ECDC. The accuracy of the author’s statistical analysis and the findings they report are not the responsibility of ECDC. ECDC is not responsible for conclusions or opinions drawn from the data provided. ECDC is not responsible for the correctness of the data and for data management, data merging and data collation after provision of the data. ECDC shall not be held liable for improper or incorrect use of the data.

## Conflict of Interest

The authors declare that they have no conflict of interest to disclose.

## Data Availability Statement

The human infection data were provided courtesy of the ECDC and are available upon request. The vector occurrence data and environmental predictor datasets used in this study are publicly available through their respective repositories. The input datasets required to run the models (presence and generated absence/background data matched with environmental predictor data, with geographical coordinates anonymised), together with the code required to reproduce all analyses and the final outputs, are available from Zenodo at https://doi.org/10.5281/zenodo.20157928 (Holle et al., 2026).

## Notes

https://doi.org/10.5281/zenodo.20157928

### Competing Interest Statement

The authors have declared no competing interest.

