## Supplementary Material for "Past and future phenology changes of zoonotic vector-borne diseases under climate and land-use change"

**Table S1: ODMAP protocol.** ODMAP (Overview, Data, Model, Assessment and Prediction) protocol outlining the main steps for data preparation and the construction of spatiotemporal species distribution models (SDMs) for the vectors *Ixodes ricinus* and *Culex pipiens*, as well as their associated viruses, Tick-borne encephalitis virus (TBEV) and West Nile virus (WNV).

| ODMAP section | ODMAP subsection | ODMAP elements | Value |
|---|---|---|---|
| Overview | Authorship | Study title | Past and future phenology changes of zoonotic vector-borne diseases under climate and land-use change |
| | | Authors | Valén Holle, Raphaëlle Klitting, Nadja Kabisch, Damaris Zurell |
| | | Contact | |
| | Model objective | Model objective | Hindcast; Forecast and transfer |
| | | Target output | Continuous habitat suitability index, Suitable vs. unsuitable habitat |
| | Focal taxon | Focal taxon | Vector models: *Ixodes ricinus* and *Culex pipiens*<br>Virus models: Tick-borne encephalitis virus (TBEV) and West Nile virus (WNV) |
| | Location | Location | Europe |
| | Scale of Analysis | Spatial extent | -31, 40, 34, 72 (xmin, xmax, ymin, ymax) |
| | | Spatial resolution | 0.5° |
| | | Temporal extent | 1970 – 2059 (1970s to 2050s) |
| | | Temporal resolution | Monthly |
| | | Boundary | Political (EU/EEA countries with the UK) |
| | Biodiversity data | Observation type | Vectors: Mixed - opportunistic and systematic professional collections<br>Viruses: Official and systematically collected infectious disease surveillance data from public health authorities |

| | | | |
|---|---|---|---|
| | | Response data type | Vectors: Presence-only (occurrence records)<br>Viruses: Presence-only (surveillance-based) |
| | Predictors | Type of Predictors | Climatic; Land use |
| | Hypotheses | Hypotheses | The distribution of vectors and viruses is mainly determined by climate and land use, either explicitly or implicitly. |
| | Assumptions | Model assumptions | Vectors and viruses are assumed to be in equilibrium with their environment; Climate and land use are the key explanatory variables for vector distribution; Climate and the occurrence probability of the corresponding vector species are the key explanatory variables for virus distribution; Observation biases for vectors and detection biases for viruses are considered negligible or adequately reduced by background/absence data generation. |
| | Algorithms | Modelling techniques | GLM (generalised linear model); GAM (generalised additive model); RF (random forest); BRT (boosted regression tree) |
| | | Model complexity | Vector models: Included a maximum of 10 predictors after variable selection – intermediate model complexity to ensure robust transfer to future environmental conditions.<br>Virus models: Included a maximum of 4 predictors after variable selection – simple to intermediate model complexity to ensure robust transfer to future environmental conditions. |
| | | Use of model averaging/ensemble modelling | The arithmetic mean was used to combine all four modelling algorithms into one ensemble. |
| | Workflow | Model workflow | Environmental data: Historical and future climate and land use data were obtained from ISIMIP and LUH2. Daily climate data were aggregated into monthly averages and totals to match the temporal resolution of the study. Future land use data from LUH2 were aggregated by a factor of two to achieve the target spatial resolution of 0.5°.<br>Vectors: Spatially-explicit vector occurrence data were obtained from GBIF and VectorMap. Occurrences were cleaned and aggregated to a 0.5° spatial and monthly temporal resolution, and duplicate records within grid cells and months of a given year were removed. Background points were generated by randomly sampling locations within a defined buffer distance from known presence locations (*Ixodes ricinus*: 150 km; *Culex* |

| | | | |
|---|---|---|---|
| | | | *pipiens:* 100 km), targeting a presence-to-background ratio of 1:10. Background points were sampled separately for each month of the year, resulting in temporally matched background data. To reduce spatial autocorrelation, all monthly presence and background points were thinned using a 50 km threshold (aiming for a checkerboard-like pattern). These points were then matched with month- and year-specific climate data and year-specific land use data. For model construction, we selected the most important and weakly correlated predictor variables. The three temperature variables (tas, tasmin, tasmax) were highly correlated but can influence species distributions differently depending on the time of the year. Therefore, we excluded them from the correlation-based variable selection process. Instead, we created three distinct predictor sets, each containing the identified most important and weakly correlated predictors along with one of the temperature variables. Spatiotemporal SDMs were fitted using four statistical algorithms, each applied to all three temperature-variant predictor sets: GLMs, GAMs, RFs, and BRTs. For regression-based methods (GLMs, GAMs), all data points were used with equal weighting of presence and background. For machine-learning methods (RFs, BRTs), *n* replicate background datasets were used at a 1:1 presence-background ratio, generating *n* models that were subsequently averaged. Ensemble models were then constructed by averaging predictions across all four algorithms, already averaged over the three temperature-specific predictor sets. Model performance was evaluated using a 5-fold cross-validation, with the focal evaluation metrics AUC and Boyce index calculated for each algorithm and their ensemble. Monthly suitability predictions for the vector species were generated for historical time frames (hindcasting: 1970-2019) and future scenarios (forecasting: 2030-2059). For hindcasting, historical factual (observed) as well as counterfactual climate and land-use simulations were included (with 1901 as the base year). For forecasting, climate data from five different models (gfdl-esm4, ipsl-cm6a-lr, mpi-esm1-2-hr, mri-esm2-0, ukesm1-0-ll) and climate and land use projections from three different SSP scenarios (SSP126, SSP370, SSP585) were used. Both continuous habitat suitability and binary presence-absence predictions (using the maxTSS threshold) were derived.<br><br>Viruses: Locally acquired, confirmed human disease case data were obtained from TESSy/ECDC at the NUTS3 level, already provided as monthly aggregated data across years. To generate spatially-explicit infection data, European NUTS3 municipalities were rasterised to the target spatial resolution of 0.5°, and the central coordinates of municipalities with reported infections were extracted. For municipalities represented by only one cell after rasterisation, these coordinates were used as the final location. Similarly, for NUTS3 municipalities too small to occupy a 0.5° cell, the central coordinates were considered as the infection point. Records stemming from large NUTS3 municipalities spanning more than one cell were excluded to minimise the spatial uncertainty of infection locations. As with |

| | | | |
|---|---|---|---|
| | | | the vector data, duplicate records reported within the same grid cell and month of a given year were removed. Absences were then randomly sampled in a balanced ratio to the presences. Absences were drawn from all cells within NUTS3 municipalities in EU/EEA countries with mandatory surveillance, selecting only those where no infection had been reported to the ECDC (also considering NUTS3 municipalities that comprised multiple cells). To ensure temporal consistency, absence points were generated separately based on disease occurrences within the same month of a given year. Spatial autocorrelation was reduced by thinning both monthly presence and absence points using a 50 km threshold (aiming for a checkerboard-like pattern). These points were then matched exclusively with year- and month-specific climate variables. Since the distribution of the studied diseases depends on the presence of their vector species, predicted monthly vector probabilities of the corresponding primary vector species were incorporated as an environmental predictor in the virus models, following a nested modelling approach. Predictor selection, model construction, validation and predictions followed the same workflow as for the vectors, with the exception that virus models were fitted using a balanced 1:1 presence-absence ratio and no predefined weighting. Therefore, the averaging of replicate sets of machine-learning models was not required. Virus models and predictions did not directly consider land use layers; however, the effects of land use change were implicitly accounted for through the incorporation of the corresponding habitat suitability predictions of the main vector species. Hindcast and forecast predictions were generated for the study area encompassing EU/EEA countries. |
| | Software | Software | R version 4.3.1 (2026-06-16 ucrt) with<br>[1] CoordinateCleaner_3.0.1<br>[2] corrplot_0.92<br>[3] countrycode_1.6.0<br>[4] dismo_1.3-14<br>[5] gbm_2.1.8.1<br>[6] giscoR_0.6.0<br>[7] ggplot2_4.0.0<br>[8] lubridate_1.9.3<br>[9] maps_3.4.1<br>[10] mgcv_1.8-42<br>[11] PresenceAbsence_1.1.11<br>[12] randomForest_4.7-1.1<br>[13] sfheaders_0.4.3<br>[14] terra_1.7-55 |

| | | | |
|---|---|---|---|
| | | | [15] tibble_3.2.1<br>[16] tidyterra_0.6.1<br>[17] tidyverse_2.0.0 |
| Data | Biodiversity data | Taxon names | Vectors: *Ixodes ricinus, Culex pipiens*<br>Viruses: Tick-borne encephalitis virus (TBEV), West Nile virus (WNV) |
| | | Taxonomic reference system | Vectors: GBIF backbone taxonomy<br>Viruses: ICTV (International Committee on Taxonomy of Viruses) |
| | | Ecological level | Vectors: Species<br>Viruses: Pathogen |
| | | Data sources | Vectors:<br>GBIF (access: January 2025; observations from 1970 to 2019 were included);<br>VectorMap (providing institutions: the Biological Records Centre, the British Museum of Natural History, the Foundation for Applied Water Research, the Smithsonian Institution, and the U.S. Army Medical Research Directorate-Georgia; access: August/September 2025; observations from 1970 to 2019 were included)<br>Viruses:<br>TESSy/ECDC (providing countries: Austria, Belgium, Bulgaria, Croatia, Cyprus, Czechia, Denmark, Estonia, Finland, France, Germany, Greece, Hungary, Italy, Latvia, Lithuania, Netherlands, Norway, Poland, Portugal, Romania, Slovakia, Slovenia, Spain and Sweden; February 2025; reported infections from 2008 to 2019 were included) |
| | | Sampling design | Vectors:<br>GBIF – opportunistic;<br>VectorMap – opportunistic<br>Viruses:<br>TESSy/ECDC – systematic (Countries participating in disease surveillance report cases on communicable diseases and related health issues) |
| | | Sample size (included in models) | Vectors: *Ixodes ricinus* (2114); *Culex pipiens* (1047)<br>Viruses: TBEV (1302); WNV (269) |

| | | | |
|---|---|---|---|
| | | Scaling | Rasterisation to 0.5°; Spatial thinning of the data to 50 km |
| | | Cleaning | Vectors: Removal of duplicate records within the same month of a given year; Removal of records with missing coordinates, equal coordinates, coordinate uncertainty > 25 km (to ensure a greater confidence that the occurrence falls within the specified cell), coordinates within a 1,000 m radius of country centroids, within a 100 m radius of biodiversity institutions; Outliers.<br>Viruses: Exclusion of records stemming from NUTS3 municipalities spanning more than one grid cell to minimise spatial uncertainty of infection locations. |
| | | Absence data | Viruses: Absence data for the viruses were generated by sampling infection-free cells within NUTS3 municipalities across EU/EEA countries with mandatory surveillance systems. Because the set of EU/EEA countries with mandatory reporting changed over time, we adjusted the list of eligible countries for each year individually, based on information from the corresponding Annual Epidemiological Reports (https://www.ecdc.europa.eu/en/publications-data/monitoring/all-annual-epidemiological-reports). Additionally, we excluded cells of NUTS3 areas from absence sampling if they had reported infections that were not considered disease occurrence points due to comprising multiple cells. Absence points were sampled in a balanced ratio to the presences and separately for each month of the year, resulting in temporally matched absence data. We treated the generated data as true absences, since they were sampled only in periods when the diseases were subject to mandatory reporting. Furthermore, the date of infection for official reporting and the NUTS3 location of infection, the key information we required to generate the absences, are mandatory reporting fields (specified in the TESSy metadata report No. 56, 21 February 2025), which increased confidence that non-reported infections represented reliable absences. |
| | | Background data | Vectors: Background data for the vector species were generated using a buffer-based approach around presence locations (*Ixodes ricinus*: 150 km; *Culex pipiens*: 100 km), while excluding the presence locations themselves. These distances reflect the differing dispersal capacities of the two species, particularly noting that tick dispersal is largely dependent on host movement, while ensuring the statistical representativeness of the background data, given the 0.5° spatial resolution. The aim was to obtain a presence-to-background ratio of 1:10 for regression-based algorithms and 1:1 for machine-learning algorithms. However, due to limitations imposed by the dispersal-adjusted buffers, the actual ratio for regression-based models was lower. Background points were sampled separately for each month of the year, resulting in temporally matched background data. |

| | | | |
|---|---|---|---|
| | Data partitioning | Training data | <u>Vectors:</u> GLMs and GAMs were fitted based on all data points; For RFs and BRTs, *n* replicate models in a presence-absence ratio of 1:1 were fitted; the value of *n* was adjusted according to the available presence-to-background ratio.<br><u>Viruses:</u> All models were fitted based on all data points after randomly sampling absences in a balanced ratio to the presences. |
| | | Validation data | A 5-fold cross-validation was used. |
| | Predictor variables | Predictor variables | 5 climatic predictors (mean near-surface air temperature, maximum near-surface air temperature, minimum near-surface air temperature, precipitation, near-surface relative humidity)<br>8 land use predictors (primary forest, primary non-forest, secondary forest, secondary non-forest, pastures, rangelands, croplands, urban areas) |
| | | Data sources | ISIMIP3a |
| | | Spatial extent | -31, 40, 34, 72 (xmin, xmax, ymin, ymax) |
| | | Spatial resolution | 0.5° |
| | | Coordinate reference system | lon/lat WGS 84 (EPSG:4326) |
| | | Temporal extent | 1970-2019 |
| | | Temporal resolution | Climate predictors: Monthly (originally daily)<br>Land-use predictors: Yearly |
| | | Data processing | Climate predictors: Daily temperature and humidity values were aggregated as monthly means, while precipitation values were aggregated as monthly totals<br>Land-use predictors: Data were aggregated into the eight distinct classes for each year |
| | Transfer data | Data sources | Climate data: ISIMIP3a (historical); ISMIP3b (future)<br>Land use data: ISIMIP3a (historical); LUH2 (future) |
| | | Spatial extent | -31, 40, 34, 72 (xmin, xmax, ymin, ymax) |

| | | | |
|---|---|---|---|
| | | Spatial resolution | Climate data: 0.5° (historical and future)<br>Land-use data: ISIMIP - 0.5°(historical); LUH2 – 0.25° (future) |
| | | Temporal extent | Hindcast: 1970-2019<br>Forecast: 2030-2059 |
| | | Temporal resolution | Climate data: Monthly (originally daily)<br>Land-use data: Yearly |
| | | Models and scenarios | Historical: Counterfactual climate and land-use data (1901 as base year)<br>Future: Five climate models (gfdl-esm4, ipsl-cm6a-lr, mpi-esm1-2-hr, mri-esm2-0, ukesm1-0-ll) and three SSPs (SSP126, SSP370, SSP585) |
| | | Data processing | Climate data: Daily temperature and humidity values were aggregated as monthly means, while precipitation values were aggregated as monthly totals.<br>Land-use data: Future rasters were aggregated by a factor of two to transition from a spatial resolution of 0.25° to the target resolution of 0.5°; Data were aggregated into the eight distinct classes. |
| | | Quantification of novelty | For environmental novelty was not checked. |
| Model | Multicollinearity | Multicollinearity | Multicollinearity was assessed, and for highly correlated pairs (Spearman $\|r\|> 0.7$), only the more important variable was retained. Variable importance was determined using explained deviance from univariate GLMs with 5-fold cross-validation. The univariate GLMs were fitted with a binomial error distribution and second-order polynomials, assigning equal weight to the sum of presences and background data for vector datasets. For the balanced presence-absence virus datasets, no predefined weights were applied. The predictive performance of each variable was then assessed through cross-validation, using the percentage of explained deviance as the evaluation metric. Because the three temperature variables (tas, tasmin, tasmax) are highly correlated but may affect species distributions differently depending on the time of the year, they were excluded from the correlation-based variable selection process. Instead, three distinct predictor sets were created, each comprising the identified most important and weakly correlated predictors along with one of the temperature variables. |

| | | | |
|---|---|---|---|
| | Model settings | Model settings (fitting) | GLM: family (binomial), formula (linear and quadratic terms), AIC-based stepwise variable selection<br>GAM: family (binomial), smoothTerms (non-parametric cubic smoothing splines (k=4))<br>RF: ntree (1000), maxnodes (10)<br>BRT: distribution (Bernoulli), interactionDepth (2), shrinkage (optimised learning rate to yield tree numbers between 1,000 and 5,000), bagFraction (0.75)<br>Vector models: GLMs and GAMs were fitted with equal weighting of presences and background data; RFs and BRTs were trained on replicated balanced datasets.<br>Virus models: No predefined weighting was applied; we ensured in the GLMs that vector suitability was always included as a linear term |
| | Model selection – model averaging – ensembles | Model averaging | Vector models: For RFs and BRTs, *n* replicate models were averaged using the arithmetic mean; For all algorithms, models were averaged across the three distinct temperature-specific predictor sets.<br>Virus models: For all algorithms, models were averaged across the three distinct temperature-specific predictor sets. |
| | | Model ensembles | Predictions over the four different algorithms were averaged using the arithmetic mean; Future predictions were additionally averaged across the five different climate models under the same SSP scenario. |
| | Threshold selection | Threshold selection | maxTSS |
| Assessment | Performance statistics | Performance on validation data | AUC; Kappa; True positive rate; True negative rate; TSS; Boyce; Explained deviance |
| | Plausibility check | Response shapes | Partial response plots of cross-validated ensemble predictions |
| Prediction | Prediction output | Prediction unit | Continuous habitat suitability ranging (0.0-1.0); Suitable vs. unsuitable habitat (0/1) |
| | Uncertainty quantification | Scenario uncertainty | Future predictions covered five different climate models and three different SSPs. |

**Table S2: Climate and land-use change projections across Europe under Shared Socioeconomic Pathway (SSP) – Representative Concentration Pathways (RCP) scenarios (2010s – 2050s).** The table summarises radiative forcing levels, underlying socioeconomic narratives, and associated implications for regional climate and land-use dynamics under SSP1-RCP2.6, SSP3-RCP7.0, and SSP5-RCP8.5. Climate outcomes include changes in mean temperature and mean annual precipitation, while land-use changes are reported as percentage shifts in major land-cover classes. All values were calculated for Europe within a defined European bounding box (31°W, 40°E, 34°N, 72°N) using datasets from ISIMIP3 (Inter-Sectoral Impact Model Intercomparison Project Phase 3; https://www.isimip.org/protocol/3/; Frieler et al., 2024) and LUH2 (Land Use Harmonization project; version 2; Hurtt et al., 2020). In addition to the reported scenario-specific values, land-use change implications are informed by Hurtt et al. (2020) and climate change implications and complemented based on findings from the AR6 (IPCC Sixth Assessment Report; IPCC, 2021).

| Scenario | Radiative forcing | Socioeconomic storyline | Implications for climate change in Europe (2010s to 2050s) | Implications for land-use change in Europe (2010s to 2050s) |
|---|---|---|---|---|
| **SSP1-RCP 2.6** | **Low emissions** | **Sustainability-focused**: Moderate population growth; Fast economic growth and technological improvements; Aiming at limiting biodiversity loss; Adoption of healthy diets with low animal-calorie shares; Early international cooperation for climate change mitigation | **+1.31°C; +17 mm/year in mean annual precipitation:**<br><br>Moderate warming across Europe (heat extremes will be primarily confined to the Mediterranean region, with a limited frequency of heat events - less than ~10 days per year); Increasing precipitation across Europe (with extreme events increasing in the northern, central, eastern, and Alpine regions) | **Regulated land-use changes:** Respected environmental boundaries; Increased bioenergy use in combination with carbon capture and storage; Reduced deforestation; Restoration of degraded forests; Controlled urban expansion<br><br>Specifically, across Europe, primary forest declines slightly (−1.47%), while secondary forest increases (+2.34%), indicating regrowth and reforestation; Agricultural land contracts, including cropland (−1.31%), pasture (−2.60%), and rangeland (−0.42%), driven by higher productivity and reduced demand; Primary open land decreases (−1.95%), while secondary open land expands (+4.89%), showing a shift toward managed and regenerating landscapes. Urban areas increase modestly (+0.52%), reflecting controlled spatial development. |

| | | | | |
|---|---|---|---|---|
| **SSP3-RCP 7.0** | **High emissions** | **Regional rivalry:** Focus on regional development; Slow economic development; Population growth is high in developing countries; High inequalities; Limited technological transfer in agriculture; Unhealthy diets dominated by animal-based products | **+2.09°C; +12 mm/year in mean annual precipitation:**<br><br>Stronger warming across Europe (heat extremes will intensify towards southern European regions, with more frequent occurrences); Slight precipitation increase across Europe (patterns will show a north–south contrast, with increases in Northern Europe and declines in the Mediterranean; Aridity will rise, particularly in the Mediterranean due to temperature increases and reduced precipitation) | **Hardly regulated land-use changes:** Expansion of cropland and pasture into forests, leading to large-scale deforestation; Urban sprawl in some regions (rather in developing than industrialised countries)<br><br>Specifically, across Europe, primary forest declines (−1.55%) while secondary forest increases only modestly (+1.83%), indicating limited restoration; Cropland decreases (−1.76%), but pasture declines only slightly (−0.81%) and rangeland increases (+0.26%), consistent with extensive, low-efficiency agriculture; Primary open land decreases (−3.11%) and secondary open land rises (+5.07%), suggesting more unmanaged or degraded landscapes rather than planned recovery; Urban growth is minimal (+0.08%). |
| **SSP5-RCP 8.5** | **Very high emissions** | **Fossil-fueled development:** High but resource-intensive economic growth; High technological progress; Strong globalisation leads to high levels of international trade; Unhealthy diets with heavy reliance on animal products; International cooperation for climate change mitigation is delayed | **+2.35°C; -3 mm/year in mean annual precipitation:**<br><br>High warming across Europe (heat extremes will intensify towards southern European regions, with high frequency - up to 60 days per year); Precipitation slightly decreases across Europe (patterns showing a north–south contrast, with increases in Northern Europe and declines in the Mediterranean; Aridity will rise, particularly in the Mediterranean due to temperature increases and reduced precipitation) | **Incompletely regulated land-use changes:** Strong expansion of cropland into pasture and forest; Slow decline in the rate of deforestation over time; More intensive urban growth<br><br>Specifically, across Europe, primary forest declines slightly (−1.61%) with only limited secondary forest recovery (+1.35%), indicating continued but slowing deforestation and weak restoration; Cropland is nearly stable (−0.03%) and pasture changes marginally (+0.06%), reflecting high yields that reduce expansion pressure; Primary open land decreases (−3.10%) while secondary open land rises modestly (+2.56%), suggesting ongoing land conversion with limited ecological recovery; Urban growth is relatively strong (+0.77%), reflecting rapid economic growth. |

**Table S3: Number of presence records considered in the spatiotemporal species distribution models (SDMs).** The table summarises the number of presence records incorporated into the spatiotemporal SDMs for the vector species *Ixodes ricinus* and *Culex pipiens*, and their associated viruses, Tick-borne encephalitis virus (TBEV) and West Nile virus (WNV). It includes the total number of records considered for each species and virus, along with the monthly distribution of these occurrences. For data sources, please see Table S1.

| | **Number of presences included in the models** | | | |
|---|---|---|---|---|
| | ***Ixodes ricinus*** | **Tick-Borne Encephalitis Virus (TBEV)** | ***Culex pipiens*** | **West Nile Virus (WNV)** |
| *All data* | 2114 | 1302 | 1047 | 269 |
| January | 18 | 21 | 18 | - |
| February | 20 | 10 | 15 | - |
| March | 105 | 10 | 22 | - |
| April | 270 | 44 | 41 | - |
| May | 428 | 131 | 102 | 1 |
| June | 408 | 224 | 191 | 4 |
| July | 287 | 239 | 138 | 45 |
| August | 232 | 181 | 202 | 102 |
| September | 166 | 147 | 172 | 89 |
| October | 100 | 139 | 93 | 25 |
| November | 58 | 107 | 33 | 1 |
| December | 22 | 49 | 20 | 2 |

**Table S4: Performance metrics of ensemble species distribution models (SDMs).** The table presents the performance results of the ensemble SDMs fitted for the primary vector species *Ixodes ricinus* and *Culex pipiens*, as well as their associated viruses, Tick-borne encephalitis virus (TBEV) and West Nile virus (WNV) in Europe. Model performance was assessed using three metrics: AUC (Area Under the Receiver Operating Characteristic Curve), the Boyce Index, and D² (Explained Deviance). All models were validated using a 5-fold cross-validation approach.

| | **Model performances** | | | | | | | | | | | |
|---|---|---|---|---|---|---|---|---|---|---|---|---|
| | ***Ixodes ricinus*** | | | **Tick-Borne Encephalitis Virus (TBEV)** | | | ***Culex pipiens*** | | | **West Nile Virus (WNV)** | | |
| Ensemble | **AUC** | **Boyce** | **D²** | **AUC** | **Boyce** | **D²** | **AUC** | **Boyce** | **D²** | **AUC** | **Boyce** | **D²** |
| *All data* | 0.73 | 0.99 | 0.45 | 0.76 | 1 | 0.18 | 0.74 | 1 | 0.39 | 0.83 | 1 | 0.27 |
| January | 0.59 | 1 | 0.38 | 0.76 | 0.93 | 0.21 | 0.65 | 0.99 | 0.35 | - | - | - |
| February | 0.69 | 0.98 | 0.43 | 0.81 | 1 | 0.19 | 0.82 | 0.70 | 0.40 | - | - | - |
| March | 0.79 | 1 | 0.47 | 0.57 | 0.58 | 0.01 | 0.79 | 0.99 | 0.41 | - | - | - |
| April | 0.76 | 1 | 0.46 | 0.68 | 1 | 0.10 | 0.69 | 0.88 | 0.37 | - | - | - |
| May | 0.70 | 1 | 0.45 | 0.74 | 0.97 | 0.14 | 0.74 | 0.92 | 0.39 | 1 | - | 0.03 |
| June | 0.74 | 0.76 | 0.46 | 0.79 | 1 | 0.20 | 0.69 | 1 | 0.38 | 0.88 | 1 | 0.23 |
| July | 0.75 | 1 | 0.46 | 0.81 | 1 | 0.24 | 0.77 | 1 | 0.41 | 0.91 | 0.99 | 0.32 |
| August | 0.74 | 1 | 0.46 | 0.85 | 1 | 0.28 | 0.73 | 1 | 0.39 | 0.92 | 0.99 | 0.35 |
| September | 0.72 | 1 | 0.44 | 0.69 | 1 | 0.09 | 0.74 | 1 | 0.40 | 0.87 | 0.89 | 0.31 |
| October | 0.69 | 1 | 0.43 | 0.68 | 1 | 0.09 | 0.79 | 1 | 0.41 | 0.77 | -0.11 | - 0.15 |
| November | 0.72 | 1 | 0.44 | 0.69 | 1 | 0.11 | 0.77 | 1 | 0.41 | 1 | - | -1.54 |
| December | 0.75 | 1 | 0.44 | 0.81 | 1 | 0.19 | 0.74 | 0.65 | 0.37 | 1 | - | -0.33 |

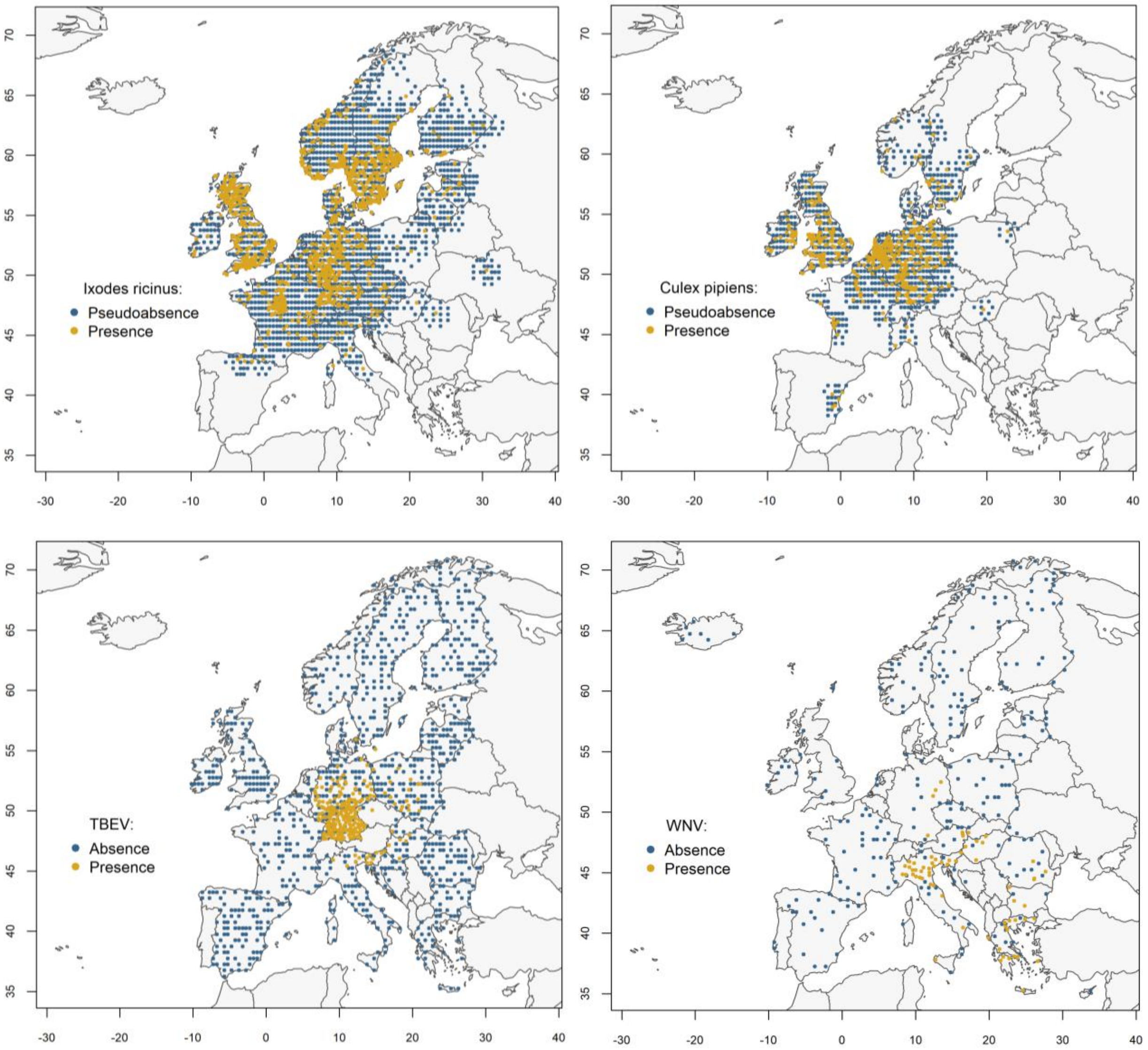


**Figure S1: Map of presence records and generated pseudoabsences/absences used in the spatiotemporal species distribution models (SDMs).** The figure displays the mapped presence records and corresponding pseudoabsence/absence locations incorporated into the SDMs for *Ixodes ricinus* (upper left), *Culex pipiens* (upper right), Tick-borne encephalitis virus (TBEV; lower left), and West Nile virus (WNV; lower right). For data sources, please see Table S1.

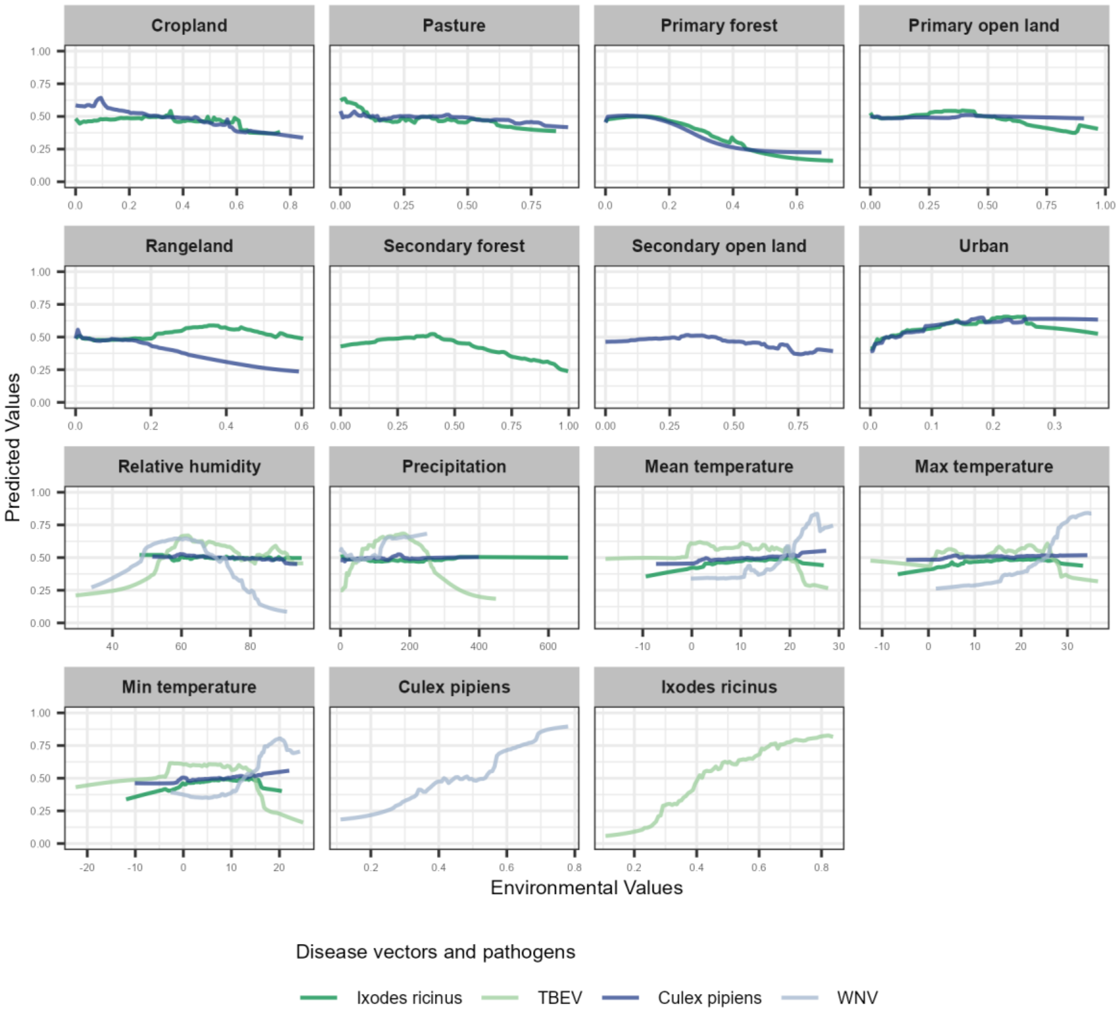


**Figure S2: Partial response curves derived from ensemble predictions across the range of environmental training data.** The figure illustrates the partial response curves for *Ixodes ricinus*, *Culex pipiens*, and their associated viruses, Tick-borne encephalitis virus (TBEV) and West Nile virus (WNV), based on ensemble model predictions to the range of environmental training data in Europe. Each curve represents the relationship between a single environmental predictor and the predicted suitability, while all other variables were held constant at their mean values. The x-axis indicates the range of each environmental variable across training data (minimum to maximum), and the y-axis shows the corresponding predicted response.

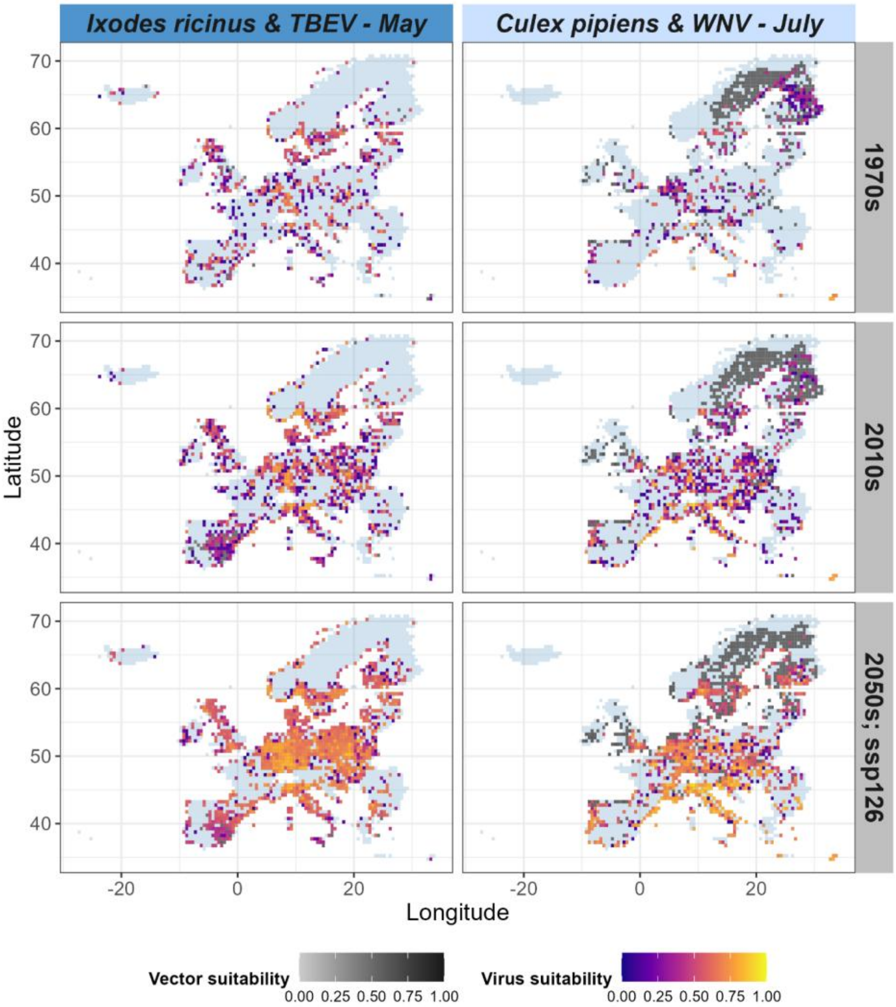


**Figure S3: Predicted distributions of vector activity and virus transmission risk over time across Europe (EU/EEA countries with UK).** Distribution trends of *Ixodes ricinus* and Tick-borne encephalitis virus (TBEV; left panels) are shown for May, while trends of *Culex pipiens* and West Nile virus (WNV; right panels) are shown for July. Risk distributions are illustrated for three target decades: 1970s, 2010s, and 2050s, with future projections based on the SSP1-2.6 scenario. Only grid cells with vector or virus suitability are displayed where the vector or the virus were predicted to be present within the selected month of a given decade. Predicted presences were determined using the threshold that maximises the True Skill Statistic (maxTSS). Predicted habitat suitability values for the vectors and viruses were averaged across each decade. Vector suitability is represented using a grey scale, with virus suitability overlaid using a coloured scale.

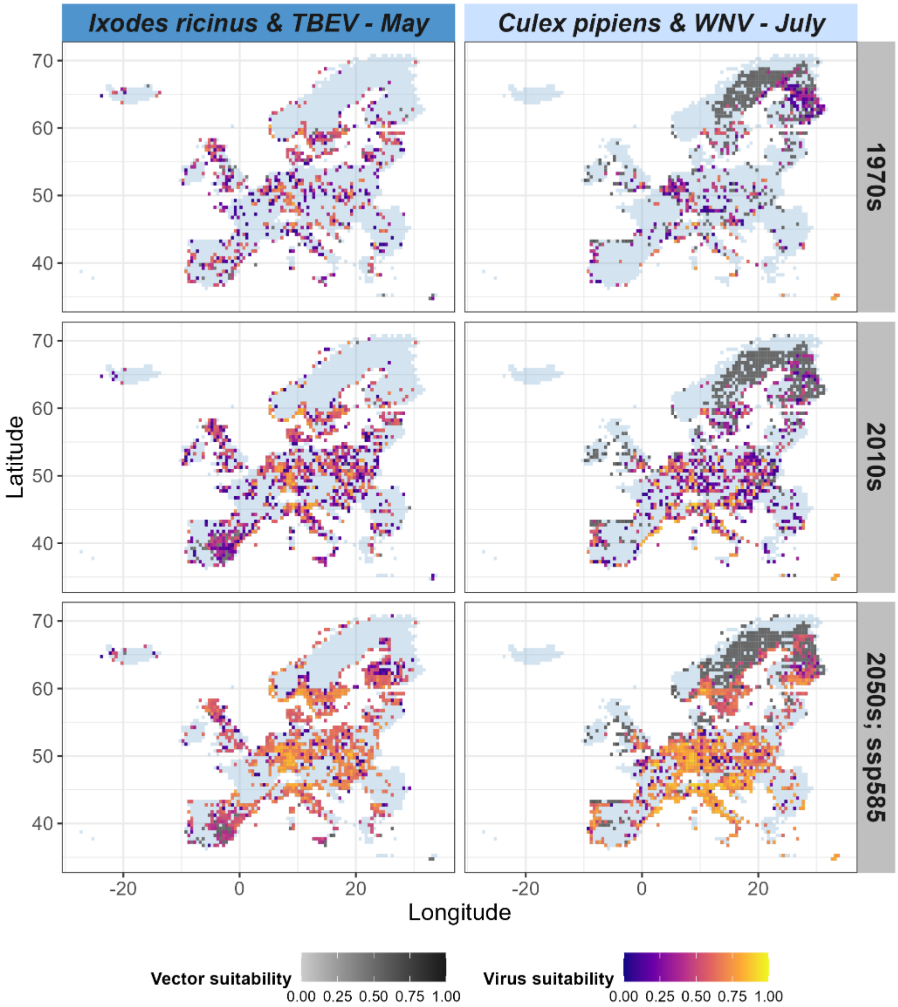


**Figure S4: Predicted distributions of vector activity and virus transmission risk over time across Europe (EU/EEA countries with UK).** Distribution trends of *Ixodes ricinus* and Tick-borne encephalitis virus (TBEV; left panels) are shown for May, while trends of *Culex pipiens* and West Nile virus (WNV; right panels) are shown for July. Risk distributions are illustrated for three target decades: 1970s, 2010s, and 2050s, with future projections based on the SSP5-8.5 scenario. Only grid cells with vector or virus suitability are displayed where the vector or the virus were predicted to be present within the selected month of a given decade. Predicted presences were determined using the threshold that maximises the True Skill Statistic (maxTSS). Predicted habitat suitability values for the vectors and viruses were averaged across each decade. Vector suitability is represented using a grey scale, with virus suitability overlaid using a coloured scale.

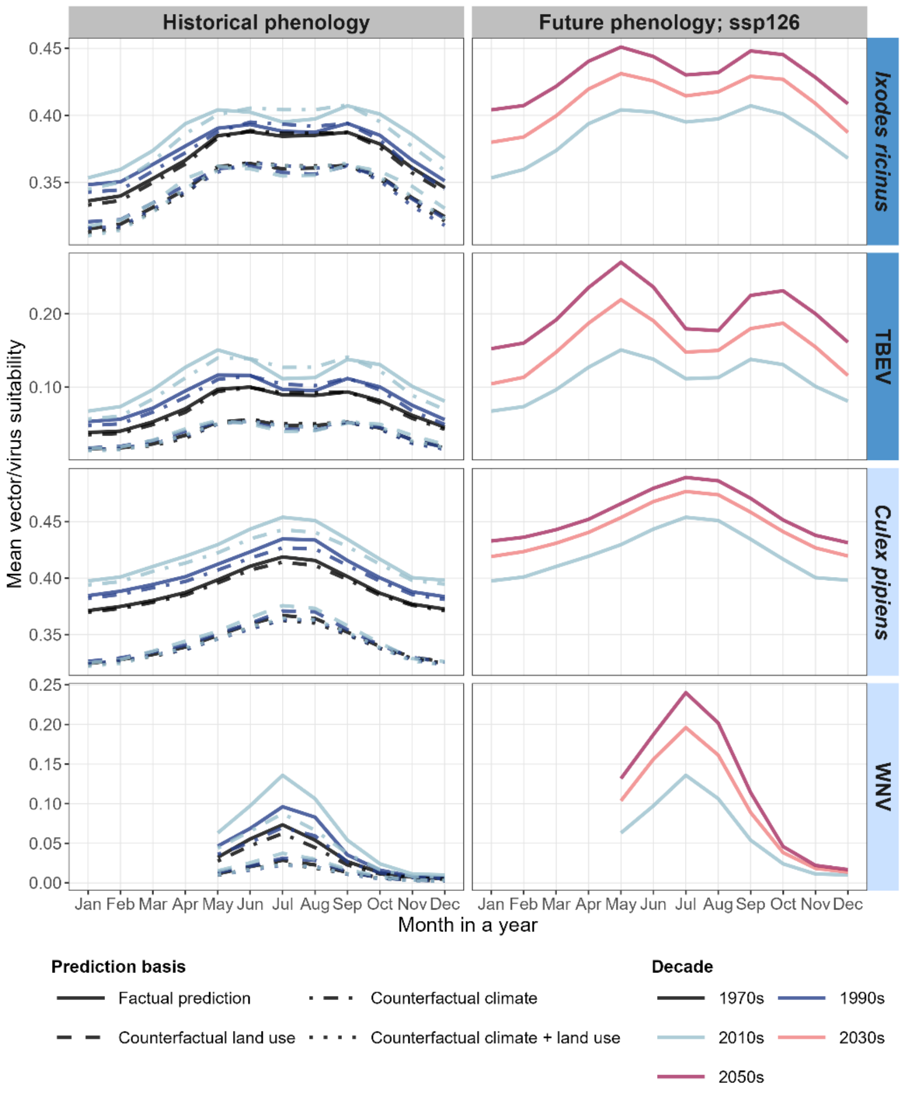


**Figure S5: Predicted seasonal risk curves of vector activity and virus transmission across historical and future decades.** Decadal trends in the seasonal phenology of activity risk for *Ixodes ricinus* and *Culex pipiens* are shown (top and third rows), along with the transmission risk for their associated viruses, Tick-borne encephalitis virus (TBEV) and West Nile virus (WNV) (second and fourth rows). Risk is represented as the average monthly predicted habitat suitability across Europe (EU/EEA countries with UK). Line colours distinguish decades (1970s, 1990s, 2010s, 2030s, 2050s), with phenologies of past decades shown in the left column and future decades in the right column. Line styles differentiate between factual and counterfactual environmental simulations (reference year 1901), including counterfactual climate, counterfactual land use, and their combination. Future projections are based on the SSP1-2.6 socio-economic scenario. For WNV, seasonal risk curves from January to April are not shown due to the absence of historical infection reports during these months.

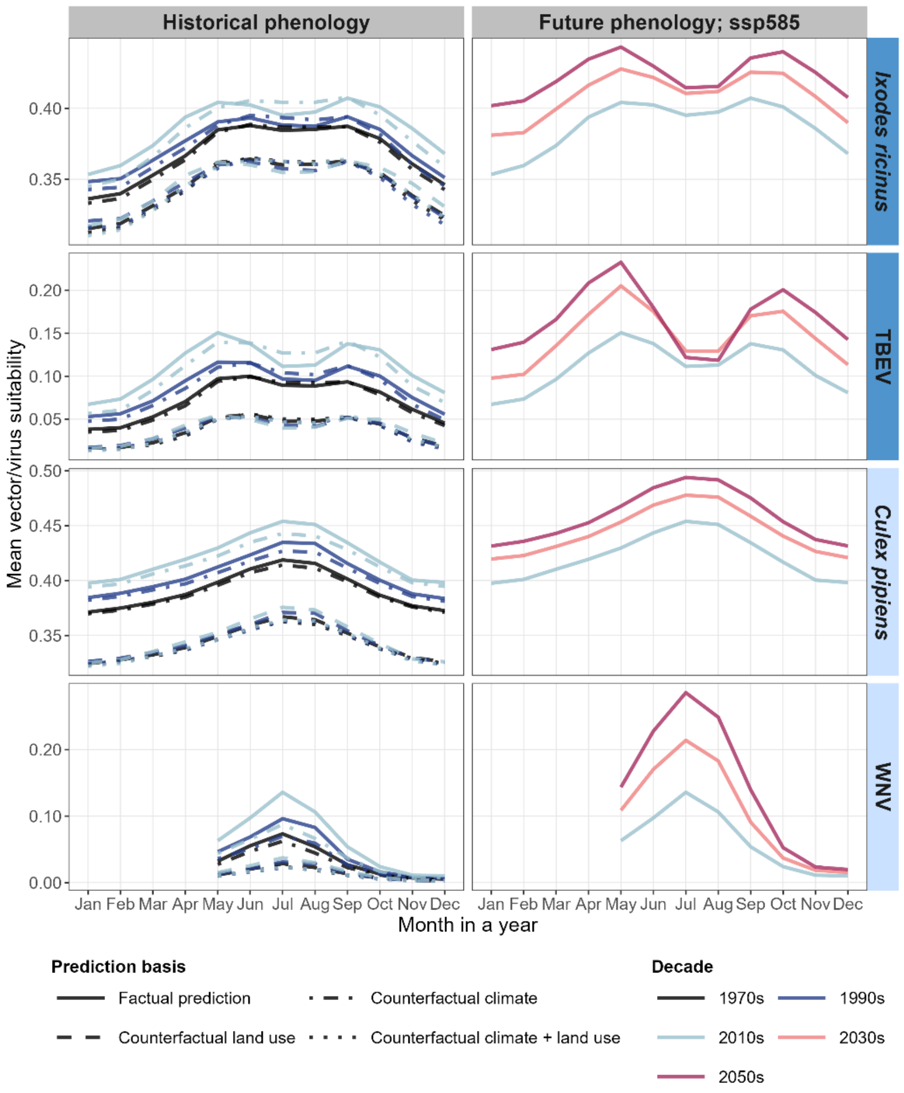


**Figure S6: Predicted seasonal risk curves of vector activity and virus transmission across historical and future decades.** Decadal trends in the seasonal phenology of activity risk for *Ixodes ricinus* and *Culex pipiens* are shown (top and third rows), along with the transmission risk for their associated viruses, Tick-borne encephalitis virus (TBEV) and West Nile virus (WNV) (second and fourth rows). Risk is represented as the average monthly predicted habitat suitability across Europe (EU/EEA countries with UK). Line colours distinguish decades (1970s, 1990s, 2010s, 2030s, 2050s), with phenologies of past decades shown in the left column and future decades in the right column. Line styles differentiate between factual and counterfactual environmental simulations (reference year 1901), including counterfactual climate, counterfactual land use, and their combination. Future projections are based on the SSP5-8.5 socio-economic scenario. For WNV, seasonal risk curves from January to April are not shown due to the absence of historical infection reports during these months.

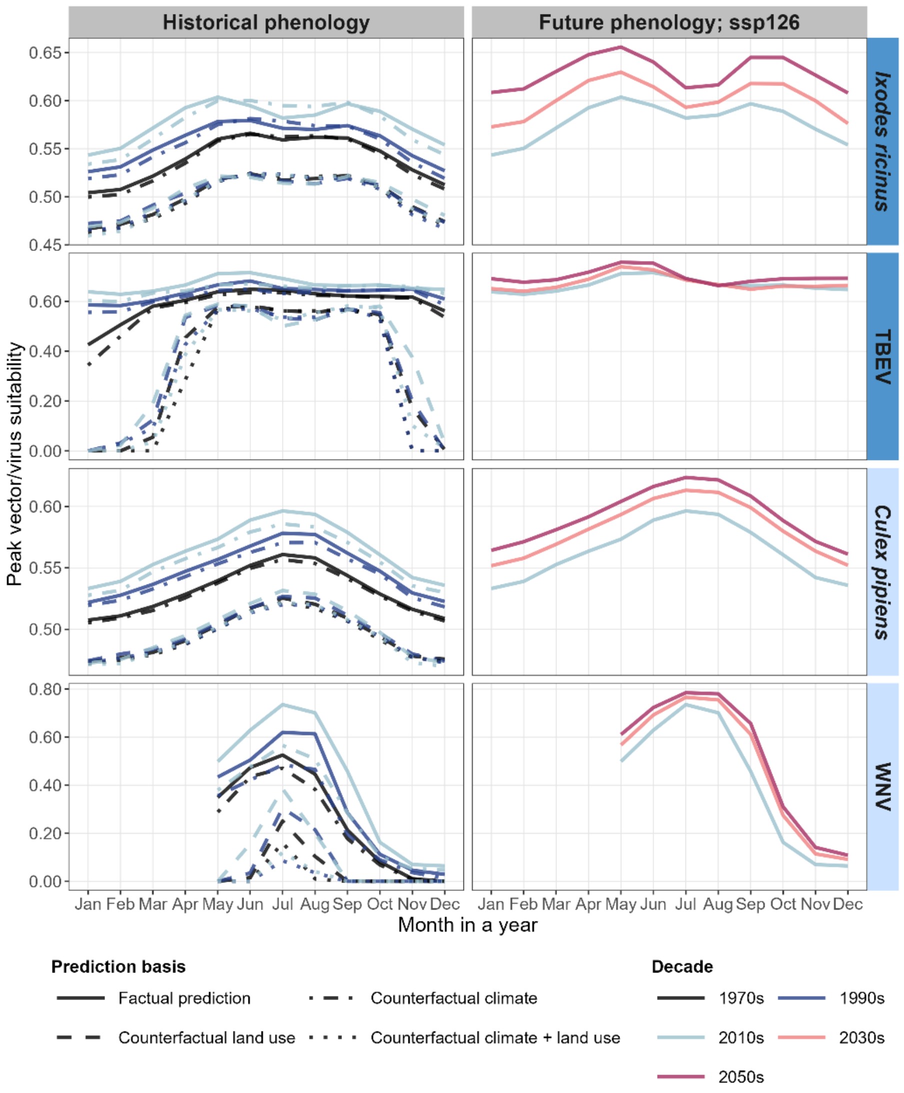


**Figure S7: Predicted seasonal risk curves of vector activity and virus transmission across historical and future decades.** Decadal trends in the seasonal phenology of activity risk for *Ixodes ricinus* and *Culex pipiens* are shown (top and third rows), along with the transmission risk for their associated viruses, Tick-borne encephalitis virus (TBEV) and West Nile virus (WNV) (second and fourth rows). Risk is represented as the peak monthly predicted habitat suitability across Europe (EU/EEA countries with UK), defined by the 95th percentile. Line colours distinguish decades (1970s, 1990s, 2010s, 2030s, 2050s), with phenologies of past decades shown in the left column and future decades in the right column. Line styles differentiate between factual and counterfactual environmental simulations (reference year 1901), including counterfactual climate, counterfactual land use, and their combination. Future projections are based on the SSP1-2.6 socio-economic scenario. For WNV, seasonal risk curves from January to April are not shown due to the absence of historical infection reports during these months.

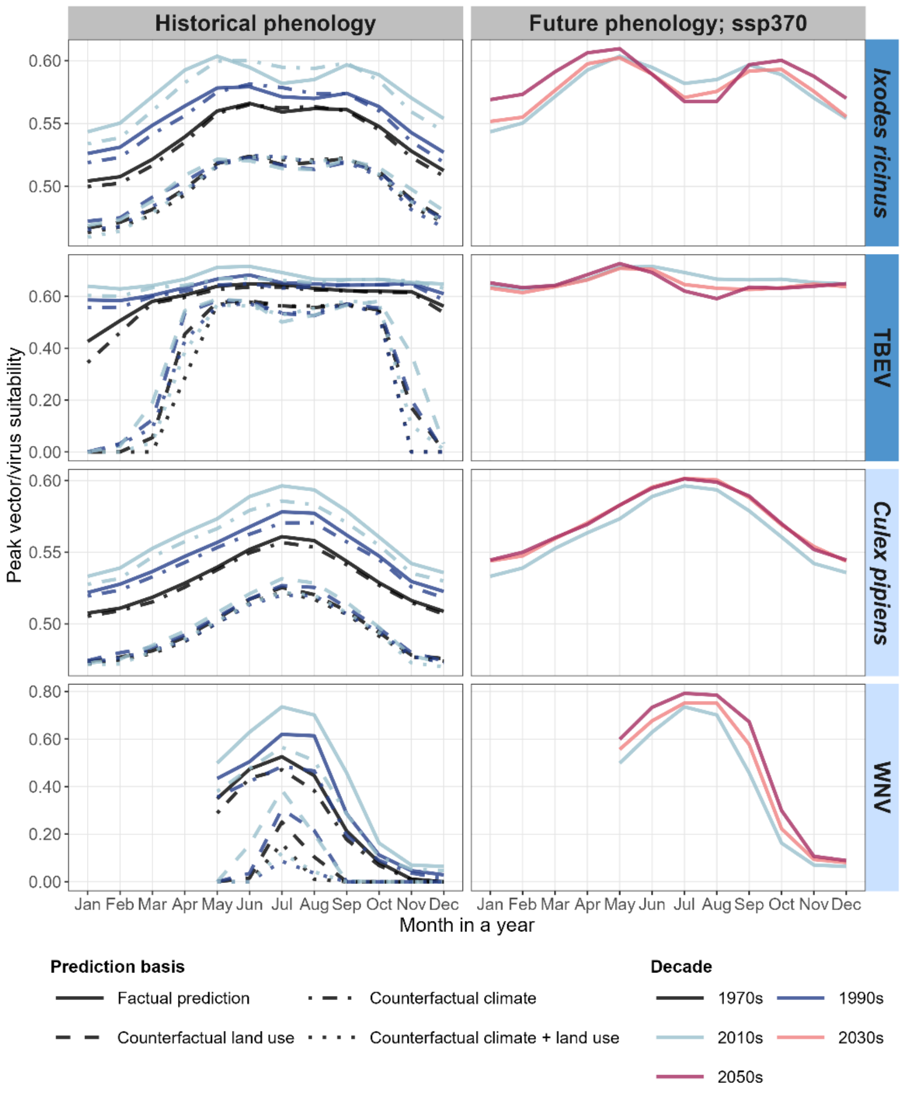


**Figure S8: Predicted seasonal risk curves of vector activity and virus transmission across historical and future decades.** Decadal trends in the seasonal phenology of activity risk for *Ixodes ricinus* and *Culex pipiens* are shown (top and third rows), along with the transmission risk for their associated viruses, Tick-borne encephalitis virus (TBEV) and West Nile virus (WNV) (second and fourth rows). Risk is represented as the peak monthly predicted habitat suitability across Europe (EU/EEA countries with UK), defined by the 95$^{th}$ percentile. Line colours distinguish decades (1970s, 1990s, 2010s, 2030s, 2050s), with phenologies of past decades shown in the left column and future decades in the right column. Line styles differentiate between factual and counterfactual environmental simulations (reference year 1901), including counterfactual climate, counterfactual land use, and their combination. Future projections are based on the SSP3-7.0 socio-economic scenario. For WNV, seasonal risk curves from January to April are not shown due to the absence of historical infection reports during these months.

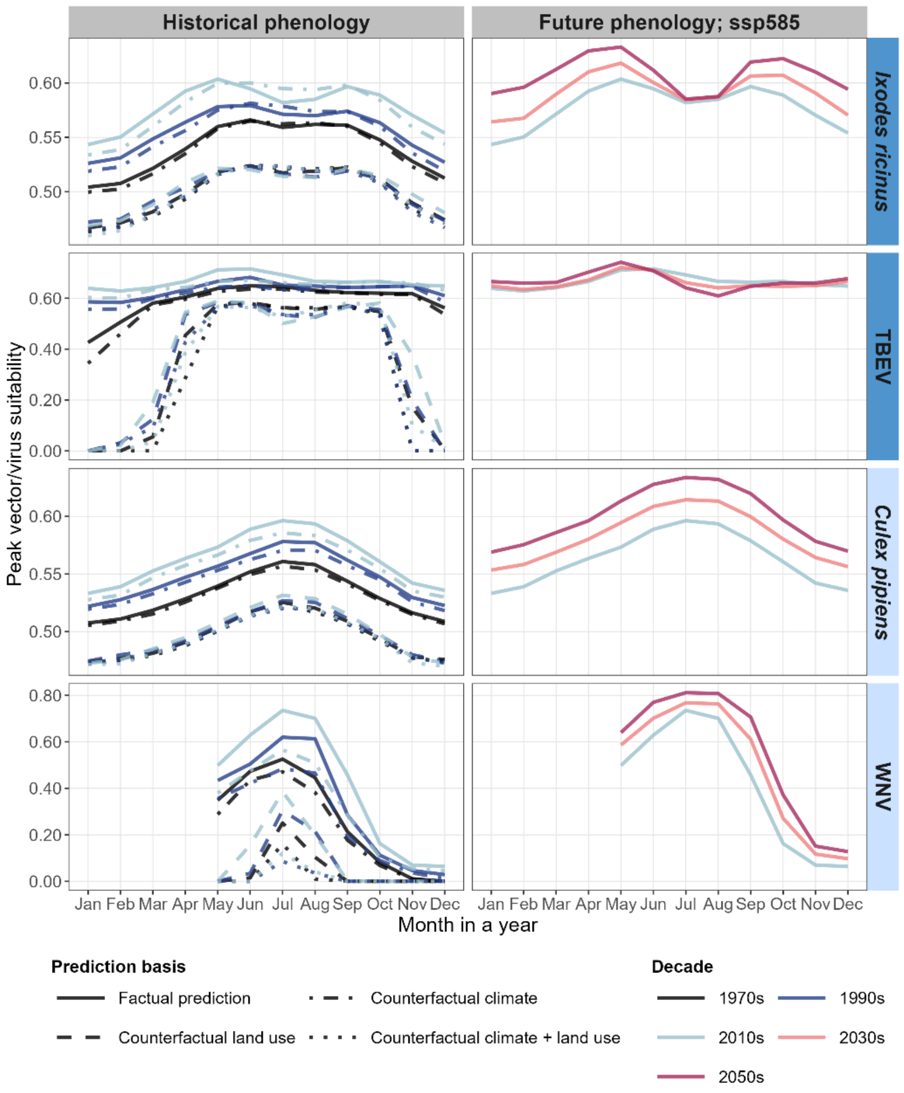


**Figure S9: Predicted seasonal risk curves of vector activity and virus transmission across historical and future decades.** Decadal trends in the seasonal phenology of activity risk for *Ixodes ricinus* and *Culex pipiens* are shown (top and third rows), along with the transmission risk for their associated viruses, Tick-borne encephalitis virus (TBEV) and West Nile virus (WNV) (second and fourth rows). Risk is represented as the peak monthly predicted habitat suitability across Europe (EU/EEA countries with UK), defined by the 95$^{th}$ percentile. Line colours distinguish decades (1970s, 1990s, 2010s, 2030s, 2050s), with phenologies of past decades shown in the left column and future decades in the right column. Line styles differentiate between factual and counterfactual environmental simulations (reference year 1901), including counterfactual climate, counterfactual land use, and their combination. Future projections are based on the SSP5-8.5 socio-economic scenario. For WNV, seasonal risk curves from January to April are not shown due to the absence of historical infection reports during these months.

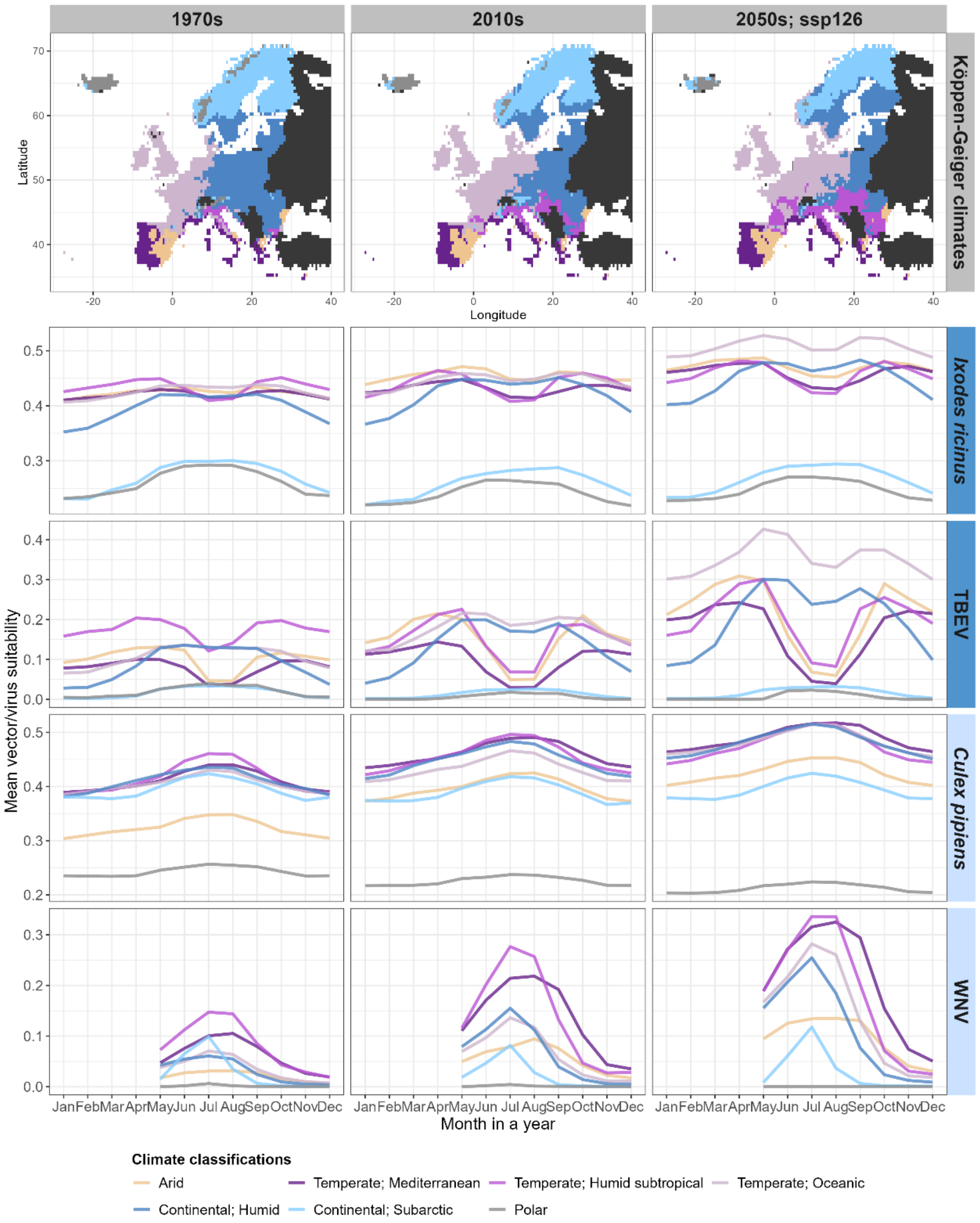


**Figure S10: Predicted seasonal risk curves of vector activity and virus transmission by Köppen-Geiger climate regions over time.** The top row shows the spatial distribution of seven different Köppen-Geiger classifications across Europe (EU/EEA countries with UK) corresponding to three time periods that align with the decades studied: 1961-1990 (representing the 1970s), 1991 – 2020 (2010s), and 2041 – 2070 (projected 2050s under the SSP1-2.6 scenario). The rows below display the seasonal phenology of activity risk *for Ixodes ricinus* and *Culex pipiens* (second and fourth rows), along with the transmission risk of their respective viruses, Tick-borne encephalitis virus (TBEV) and West Nile virus (WNV) (third and fifth rows). Vector activity and transmission probability are expressed as the average monthly predicted habitat suitability across the prevailing Köppen-Geiger climate zones in each respective time period. Line colours indicate the different Köppen-Geiger classifications: Arid, Temperate (Mediterranean, Humid subtropical, Oceanic), Continental (Humid, Subarctic), and Polar. For WNV, seasonal risk curves from January to April are not shown due to the absence of historical infection reports during these months.

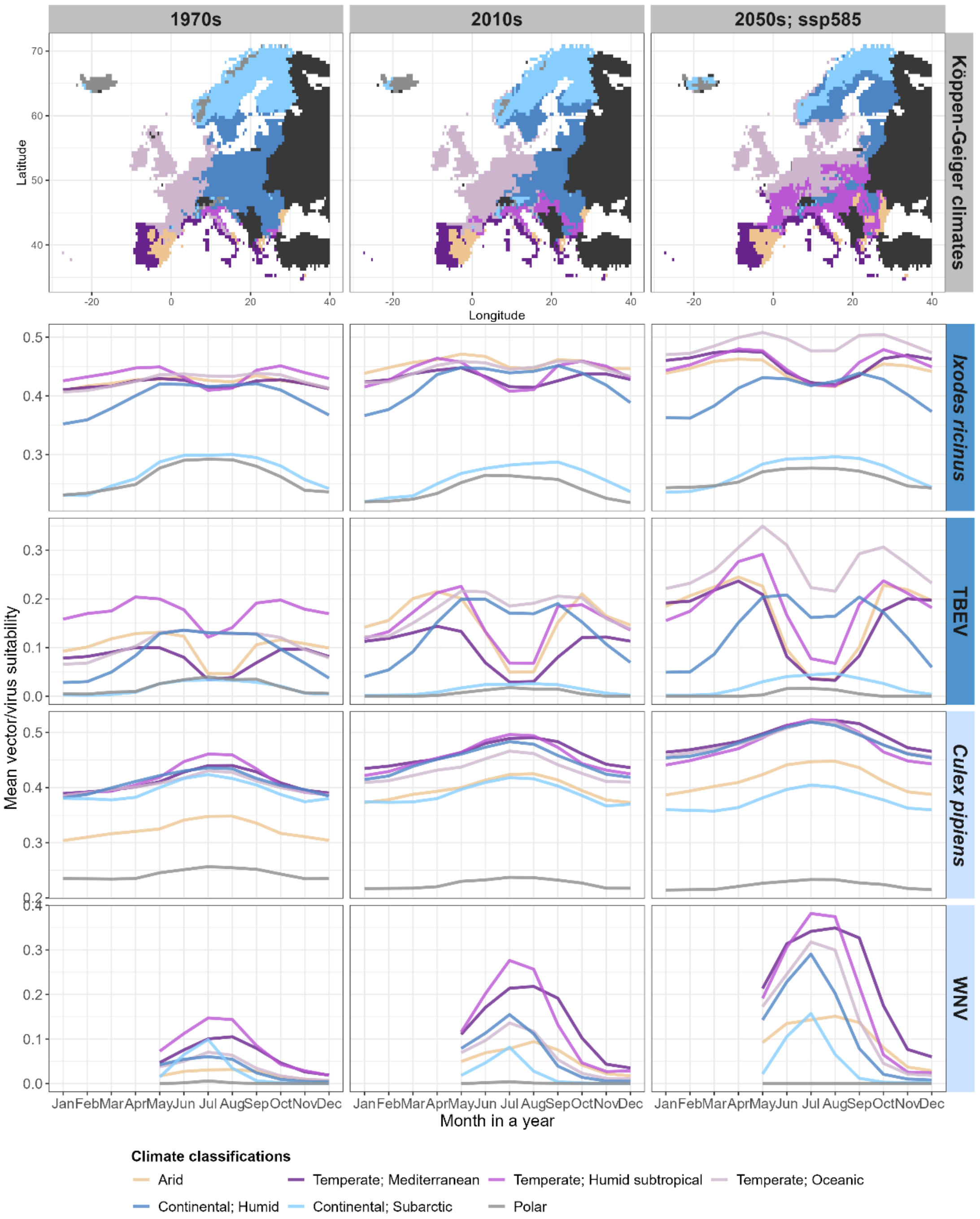


**Figure S11: Predicted seasonal risk curves of vector activity and virus transmission by Köppen-Geiger climate regions over time.** The top row shows the spatial distribution of seven different Köppen-Geiger classifications across Europe (EU/EEA countries with UK) corresponding to three time periods that align with the decades studied: 1961-1990 (representing the 1970s), 1991 – 2020 (2010s), and 2041 – 2070 (projected 2050s under the SSP5-8.5 scenario). The rows below display the seasonal phenology of activity risk for *Ixodes ricinus* and *Culex pipiens* (second and fourth rows), along with the transmission risk of their respective viruses, Tick-borne encephalitis virus (TBEV) and West Nile virus (WNV) (third and fifth rows). Vector activity and transmission probability are expressed as the average monthly predicted habitat suitability across the prevailing Köppen-Geiger climate zones in each respective time period. Line colours indicate the different Köppen-Geiger classifications: Arid, Temperate (Mediterranean, Humid subtropical, Oceanic), Continental (Humid, Subarctic), and Polar. For WNV, seasonal risk curves from January to April are not shown due to the absence of historical infection reports during these months.

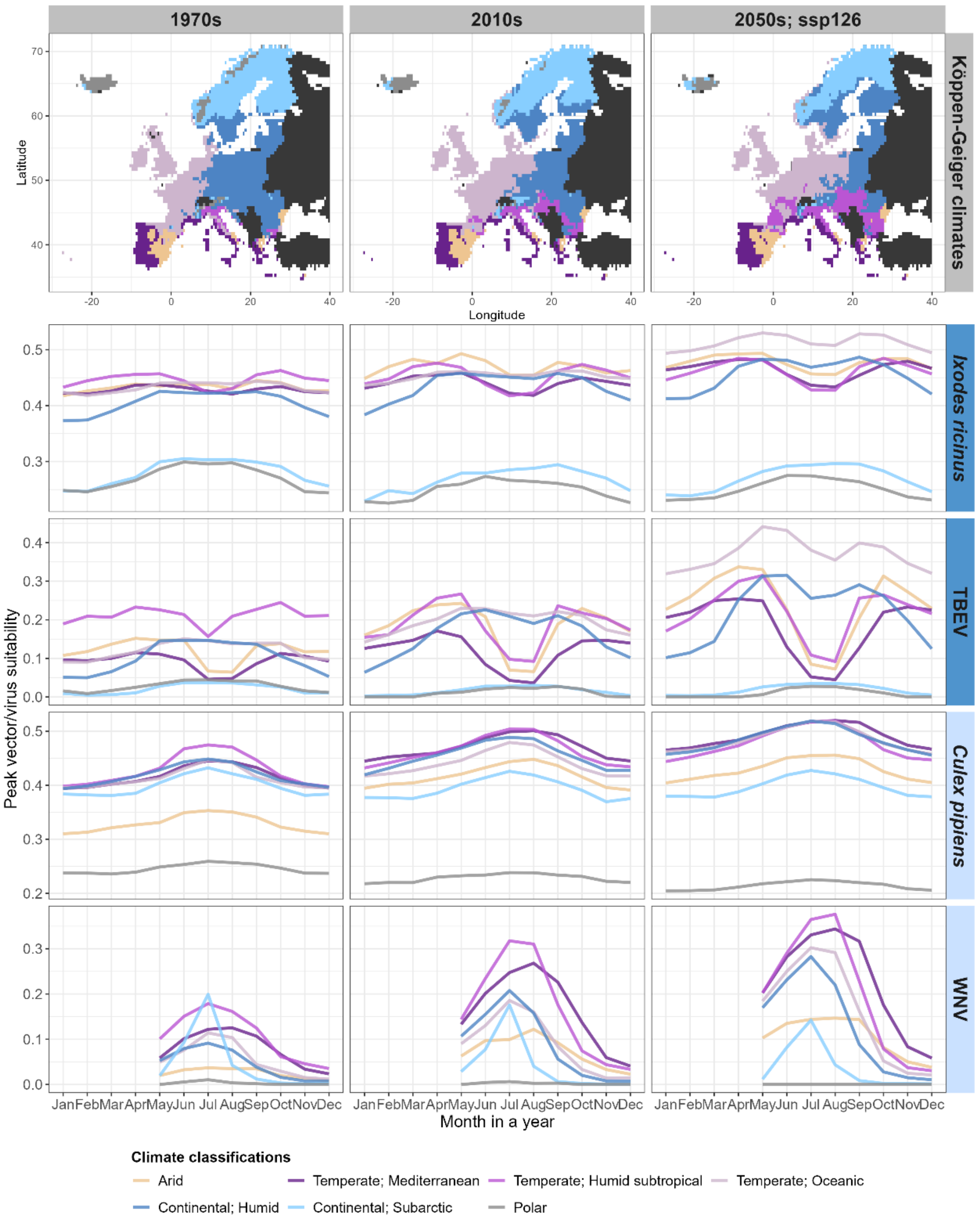


**Figure S12: Predicted seasonal risk curves of vector activity and virus transmission by Köppen-Geiger climate regions over time.** The top row shows the spatial distribution of seven different Köppen-Geiger classifications across Europe (EU/EEA countries with UK) corresponding to three time periods that align with the decades studied: 1961-1990 (representing the 1970s), 1991 – 2020 (2010s), and 2041 – 2070 (projected 2050s under the SSP1-2.6 scenario). The rows below display the seasonal phenology of activity risk for *Ixodes ricinus* and *Culex pipiens* (second and fourth rows), along with the transmission risk of their respective viruses, Tick-borne encephalitis virus (TBEV) and West Nile virus (WNV) (third and fifth rows). Vector activity and transmission probability are expressed as the peak monthly predicted habitat suitability, defined by the 95th percentile, across the prevailing Köppen-Geiger climate zones in each respective time period. Line colours indicate the different Köppen-Geiger classifications: Arid, Temperate (Mediterranean, Humid subtropical, Oceanic), Continental (Humid, Subarctic), and Polar. For WNV, seasonal risk curves from January to April are not shown due to the absence of historical infection reports during these months.

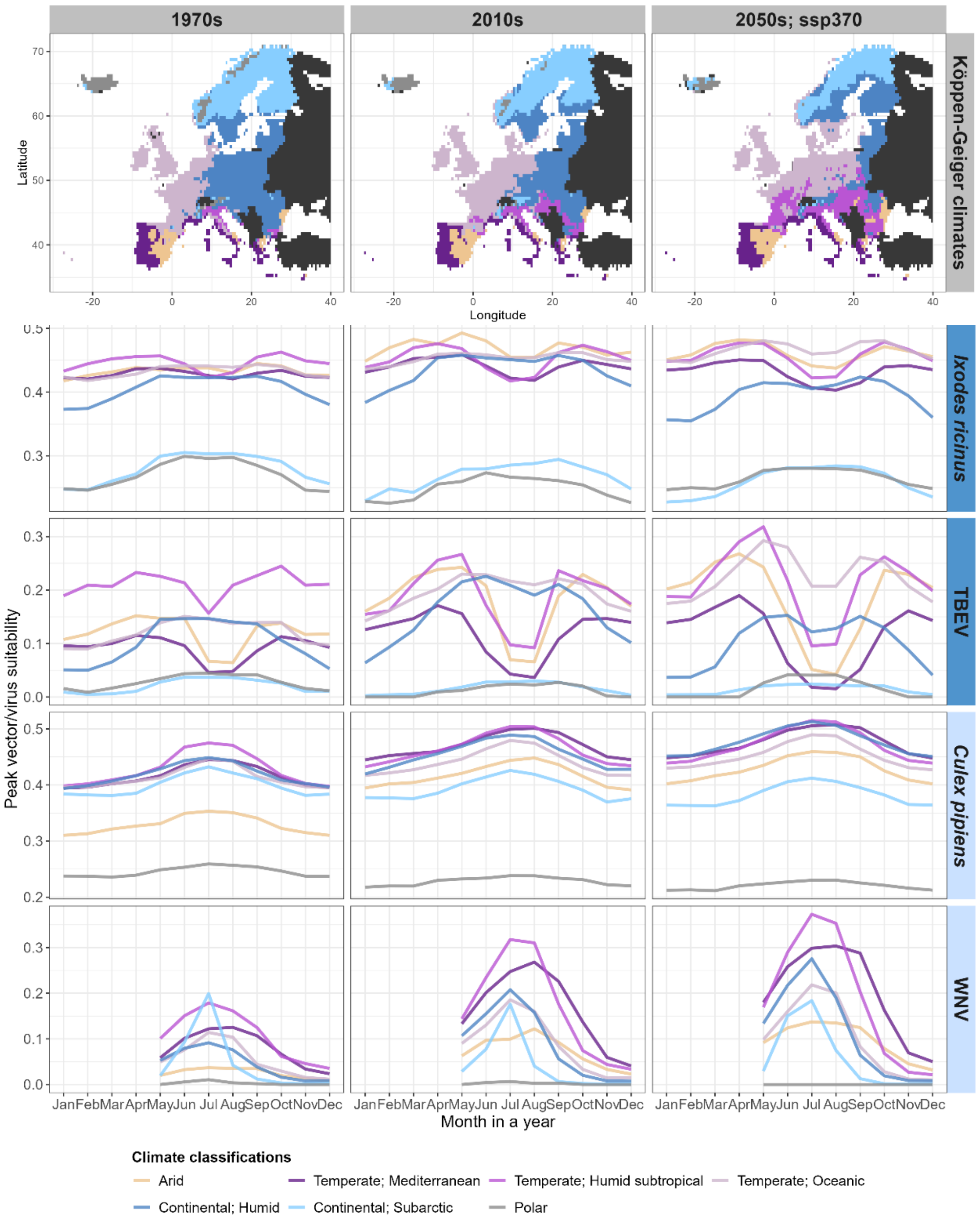


**Figure S13: Predicted seasonal risk curves of vector activity and virus transmission by Köppen-Geiger climate regions over time.** The top row shows the spatial distribution of seven different Köppen-Geiger classifications across Europe (EU/EEA countries with UK) corresponding to three time periods that align with the decades studied: 1961-1990 (representing the 1970s), 1991 – 2020 (2010s), and 2041 – 2070 (projected 2050s under the SSP3-7.0 scenario). The rows below display the seasonal phenology of activity risk for *Ixodes ricinus* and *Culex pipiens* (second and fourth rows), along with the transmission risk of their respective viruses, Tick-borne encephalitis virus (TBEV) and West Nile virus (WNV) (third and fifth rows). Vector activity and transmission probability are expressed as the peak monthly predicted habitat suitability, defined by the 95$^{th}$ percentile, across the prevailing Köppen-Geiger climate zones in each respective time period. Line colours indicate the different Köppen-Geiger classifications: Arid, Temperate (Mediterranean, Humid subtropical, Oceanic), Continental (Humid, Subarctic), and Polar. For WNV, seasonal risk curves from January to April are not shown due to the absence of historical infection reports during these months.

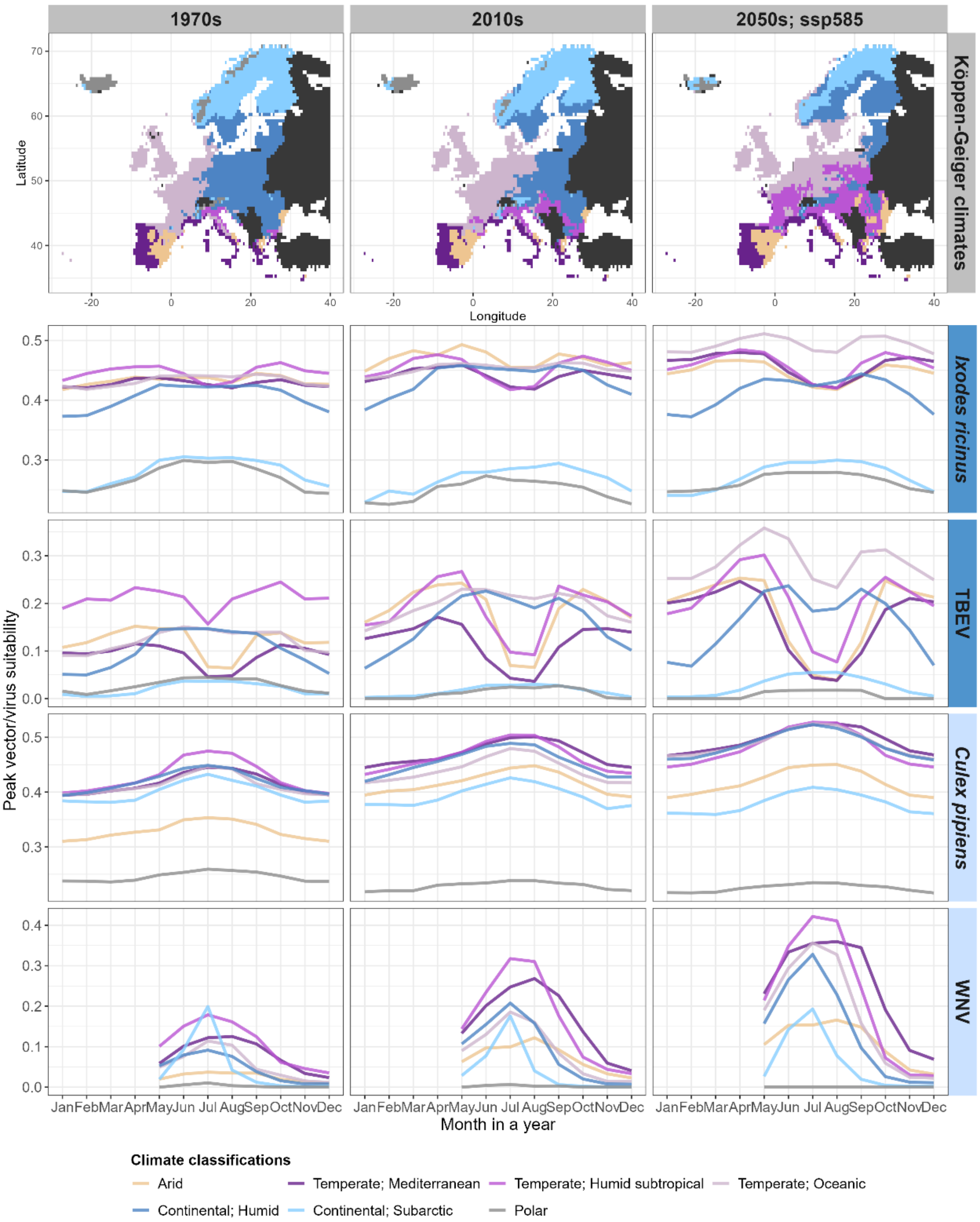


**Figure S14: Predicted seasonal risk curves of vector activity and virus transmission by Köppen-Geiger climate regions over time.** The top row shows the spatial distribution of seven different Köppen-Geiger classifications across Europe (EU/EEA countries with UK) corresponding to three time periods that align with the decades studied: 1961-1990 (representing the 1970s), 1991 – 2020 (2010s), and 2041 – 2070 (projected 2050s under the SSP5-8.5 scenario). The rows below display the seasonal phenology of activity risk for *Ixodes ricinus* and *Culex pipiens* (second and fourth rows), along with the transmission risk of their respective viruses, Tick-borne encephalitis virus (TBEV) and West Nile virus (WNV) (third and fifth rows). Vector activity and transmission probability are expressed as the peak monthly predicted habitat suitability, defined by the 95$^{th}$ percentile, across the prevailing Köppen-Geiger climate zones in each respective time period. Line colours indicate the different Köppen-Geiger classifications: Arid, Temperate (Mediterranean, Humid subtropical, Oceanic), Continental (Humid, Subarctic), and Polar. For WNV, seasonal risk curves from January to April are not shown due to the absence of historical infection reports during these months.

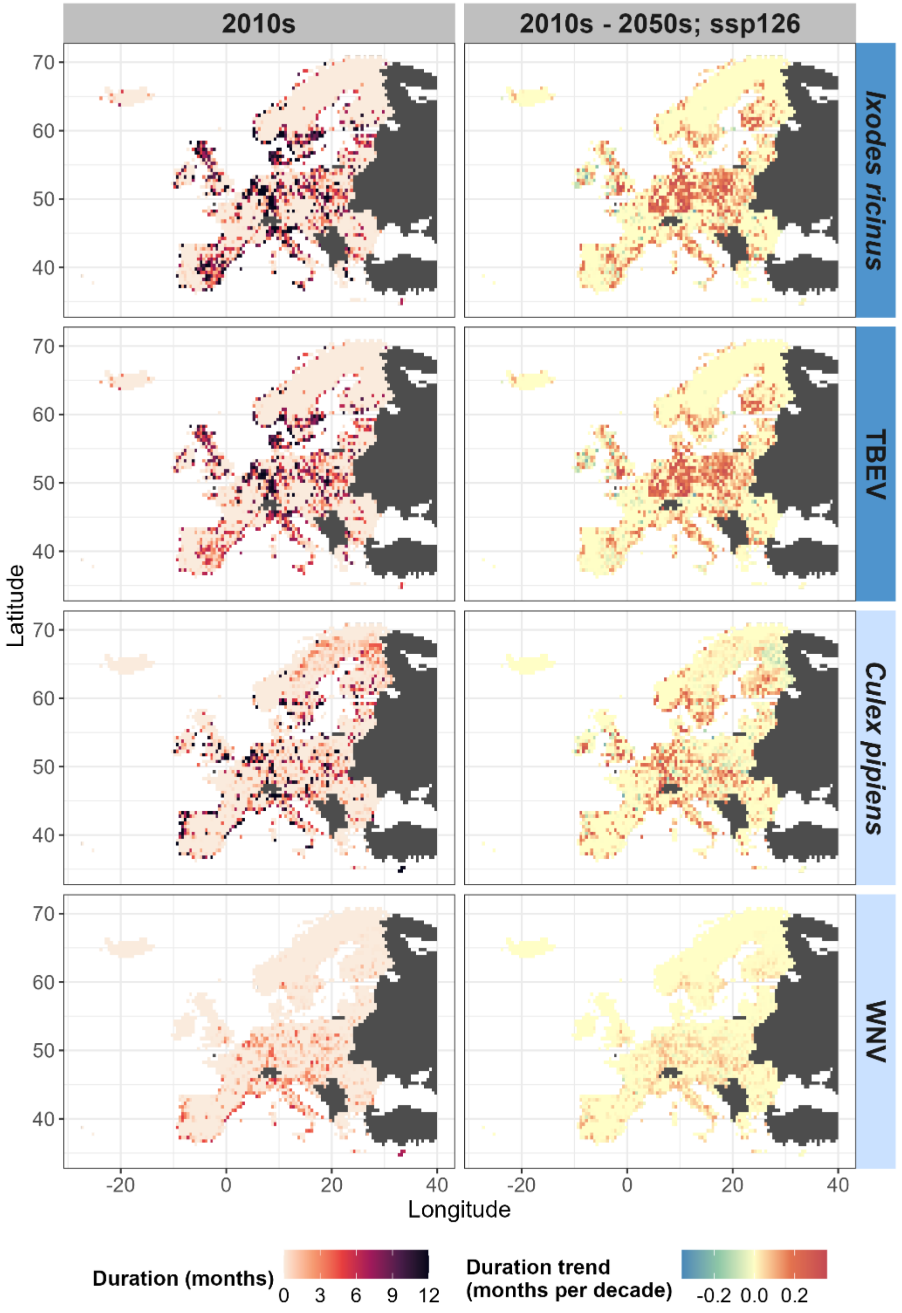


**Figure S15: Predicted duration and temporal trends of seasonal risk periods for vector activity and virus transmission across Europe (EU/EEA countries with UK).** The left column maps the average duration (in months) of seasonal risk during the 2010s for *Ixodes ricinus* and *Culex pipiens* (first and third rows), and their associated viruses, Tick-borne encephalitis virus (TBEV) and West Nile virus (WNV) (second and fourth rows). Darker shades in the left column indicate longer risk periods, up to 12 months. The right column shows the respective projected trends in seasonal risk duration for each species and virus presented in the left column, from the 2010s to the 2050s under the SSP1-2.6 scenario, expressed as the rate of change in months per decade. Warmer (yellow-red) and cooler (blue-green) tones in the right column represent projected increases or decreases in risk duration, respectively. Seasonal risk durations are derived from binary maps of monthly predicted habitat suitability, thresholded to maximise the True Skill Statistic (maxTSS).

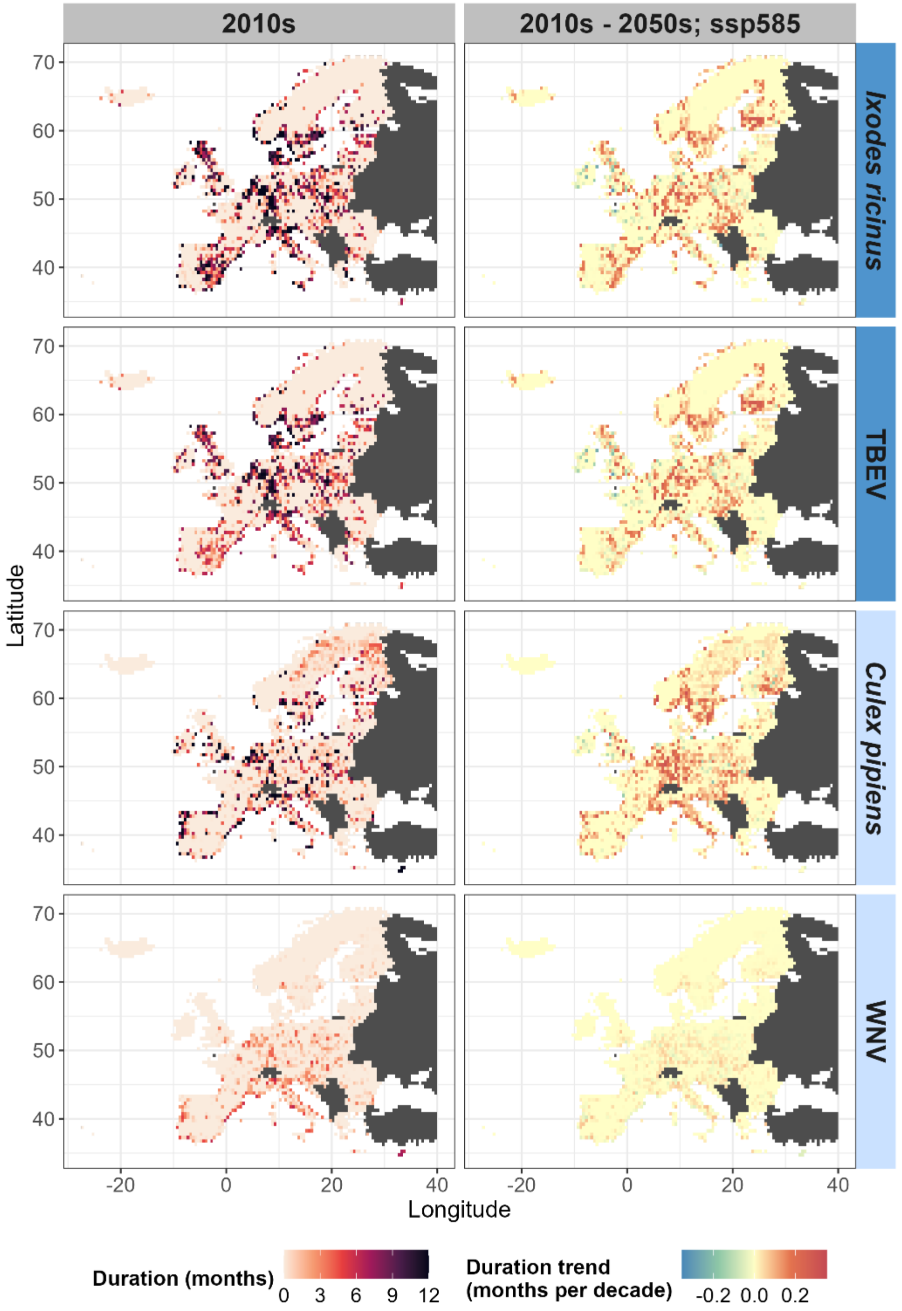


**Figure S16: Predicted duration and temporal trends of seasonal risk periods for vector activity and virus transmission across Europe (EU/EEA countries with UK).** The left column maps the average duration (in months) of seasonal risk during the 2010s for *Ixodes ricinus* and *Culex pipiens* (first and third rows), and their associated viruses, Tick-borne encephalitis virus (TBEV) and West Nile virus (WNV) (second and fourth rows). Darker shades in the left column indicate longer risk periods, up to 12 months. The right column shows the respective projected trends in seasonal risk duration for each species and virus presented in the left column, from the 2010s to the 2050s under the SSP5-8.5 scenario, expressed as the rate of change in months per decade. Warmer (yellow-red) and cooler (blue-green) tones in the right column represent projected increases or decreases in risk duration, respectively. Seasonal risk durations are derived from binary maps of monthly predicted habitat suitability, thresholded to maximise the True Skill Statistic (maxTSS).
